# AP-3 complex sorts preferential cargo to govern dense core vesicle function in neuroendocrine cells

**DOI:** 10.1101/2025.06.27.661901

**Authors:** Shashank Saxena, Vinayak Ghosh, Chandramouli Mukherjee, Badal Singh Chauhan, Sanchari Banerjee, Bhavani Shankar Sahu

## Abstract

This study reveals new insights into the role of the Adaptor protein (AP-3) complex in dense core vesicle function. Despite numerous studies, an existing knowledge lacuna in the role of AP-3 in DCV function prompted us to delve deeper. Advanced microscopy and biochemical analysis revealed compromised DCV exocytosis in AP-3-depleted PC12 cells and C. *elegans*. AP-3 depletion altered the size and positioning of DCVs. Golgi defects and RUSH (Retention under Selective Hook) substantiated the role of AP-3 in trans-Golgi DCV budding. Proteomics revealed the loss of specific known and putative novel DCV proteins, which were mislocalized and rerouted to lysosomes in AP-3-depleted cells. Bioinformatics, Proximity ligation assays and Co-immunoprecipitation identified interactions of mislocalized proteins with AP-3 subunit. These findings corroborated with functional defects in granule maturation, release modes, Zinc and neurotransmitter mobilisation. Our study highlights the complexity of the AP-3 complex in regulating DCV function and its importance in vesicle transport in neurons and neuroendocrine cells.

**Summary:** This work reveals the critical role of AP-3 complex in DCV function, highlighting its impact on DCV exocytosis, positioning, and trans-Golgi budding. This study identifies Dlk1 as a novel DCV cargo.

## Introduction

Adaptor protein (AP) complexes are heterotetrameric coat protein complexes that dictate the loading of membrane lipids, membrane proteins, and the components for the vesicular formation and fusion at the Golgi, Endosomes or at the Plasma Membrane^1–3^. APs are paralogous groups of proteins composed of distinct subunits constituting five different APs (AP1-AP5)^4–7^. AP-3 consists of two larger subunits identified as β and δ and two smaller subunits identified as µ and σ, where the δ subunit is unique to AP-3 complex^8^. The β subunit of AP-3 has ubiquitously expressed (A) and neuron-specific (B) isoforms, where the neuron-specific form is crucial for the biogenesis and release of DCVs and synaptic vesicles^9,10^. APs interact with cargo proteins involve tyrosine-based sorting motifs (YXXΦ; where X can be polar residues and Φ is a bulky hydrophobic residue)^11^ as well as di-leucine motifs ([DE]xxxL[LI]; consisting of acidic residues, where X can be any amino acid) at the cytoplasmic tail^4^. AP-3 complex is recruited to the TGN and endosomal compartments^12^ with the help of ADP-Ribosylation factor 1 (ARF1)^13^ participating in the transport of Cargo towards the lysosome^14^ and lysosome-related organelles such as melanosomes and platelets^15,16^. Work-related to AP-3 has led to the development of two mouse knockout models lacking either the δ subunit(Mocha)^17^ or β subunit(Pearl)^18^. The mocha mice lack the δ adaptin and, therefore, lack functional AP-3^17^. While the Pearl mice have a scarcity of ubiquitously expressing A isoform of the β subunit, the Brain-specific B isoform is unaltered^18^. AP-3 deficiency in mocha mice does impair stimulus-coupled DCV release in adrenal chromaffin cells, increases quantal size, and overexpression of AP-3 reverses this effect^19,20^. While previous reports indicate defective biogenesis and impaired exocytosis of DCVs in the absence of AP-3^19–21^, the precise mechanisms through which AP-3 influences specific Cargo and regulates other DCV functions is intriguing and worth further investigation.

We have used a multidisciplinary approach to address these hitherto understudied aspects of the AP3 complex in regulating DCV function. First, we disrupted the AP-3 function by targeting its core δ subunit using multiple shRNAs^22^. Functional analyses of DCVs, including microscopy and biochemical assays, revealed compromised DCV exocytosis in AP-3 depleted cells. Ultrastructural transmission electron microscopy (TEM) and morphometric analysis further showed an increased proportion of immature DCVs positioned farther from the plasma membrane. This finding was consistent with enhanced co-localization of DCVs with syntaxin 6, a marker for immature DCVs that is typically removed during maturation^23^. To broaden our understanding, we conducted comparative quantitative organellar proteomics on vesicle-enriched fractions, as described previously ^24^. Our analysis demonstrated the depletion of key membrane proteins associated with DCVs, including known markers (Syt1, VMAT1, ZnT3)^25–27^ and putative novel candidates (DLK-1, SCN3β). Confocal microscopy revealed mislocalization of these proteins to lysosomes rather than DCVs, suggesting that AP-3 deficiency leads to missorting and rerouting of DCV cargo to lysosomes. Bioinformatics analysis identified binding motifs in these candidate proteins for subunits of the AP-3 complex, further supporting their dependence on AP-3 for correct sorting.

Functionally, AP-3 depletion resulted in compromised DCV compartments, as evidenced by reduced neurotransmitter and zinc mobilization—hallmarks of mature DCVs^28,29^. These functional defects were attributed to the mislocalization of VMAT1 and ZnT3. Additionally, we observed a significant reduction in full-fusion exocytosis events, likely due to the missorting of Syt-1, a key protein in exocytotic fusion pore formation. In summary, our study demonstrates that the AP-3 complex plays a critical and multifaceted role in sorting proteins to the DCV compartment. Future studies should explore its interactions with other adaptor protein complexes, such as AP-1, which has been implicated in DCV biogenesis^30,31^. Moreover, the specific cargo sorting pathways of the AP-3 complex—whether from the trans-Golgi network or recycling endosomes to DCVs—require further investigation, which we plan to address in future work.

## Results

### *apb-3* null C.elegans has impaired NLP-21 exocytosis from DNC to coelomocytes

In C.elegans, AP-3 is expressed from a single gene having β, δ, µ and σ subunits^32^. Here, we used an AP-3 null worm with a mutant β subunit (*apb-3*) to study the dynamics of DCV^33^. DCV cargo secretion could be studied by measuring the levels of NLP-21(neuropeptide-like protein 21, neuropeptide analogue of C.elegans) in coelomocytes, scavenger cells which take up the NLP-21 secreted from the DCVs of the Dorsal Nerve Cord (DNC)^34–36^(Fig 1A). We tagged NLP-21 with YFP to detect its release from the DNC and uptake by coelomocytes. Our findings revealed an impairment in exocytosis of NLP-21 from the DCVs in DNC, as evidenced by an increased abundance of NLP-21 in the DNC of *apb-3* null worms(Fig 1B-C). Correspondingly, a reduced abundance of NLP-21 in coelomocytes compared to their wild-type counterpart (Fig 1D-F) signifies an impaired DCV exocytosis.

**Figure 1.**
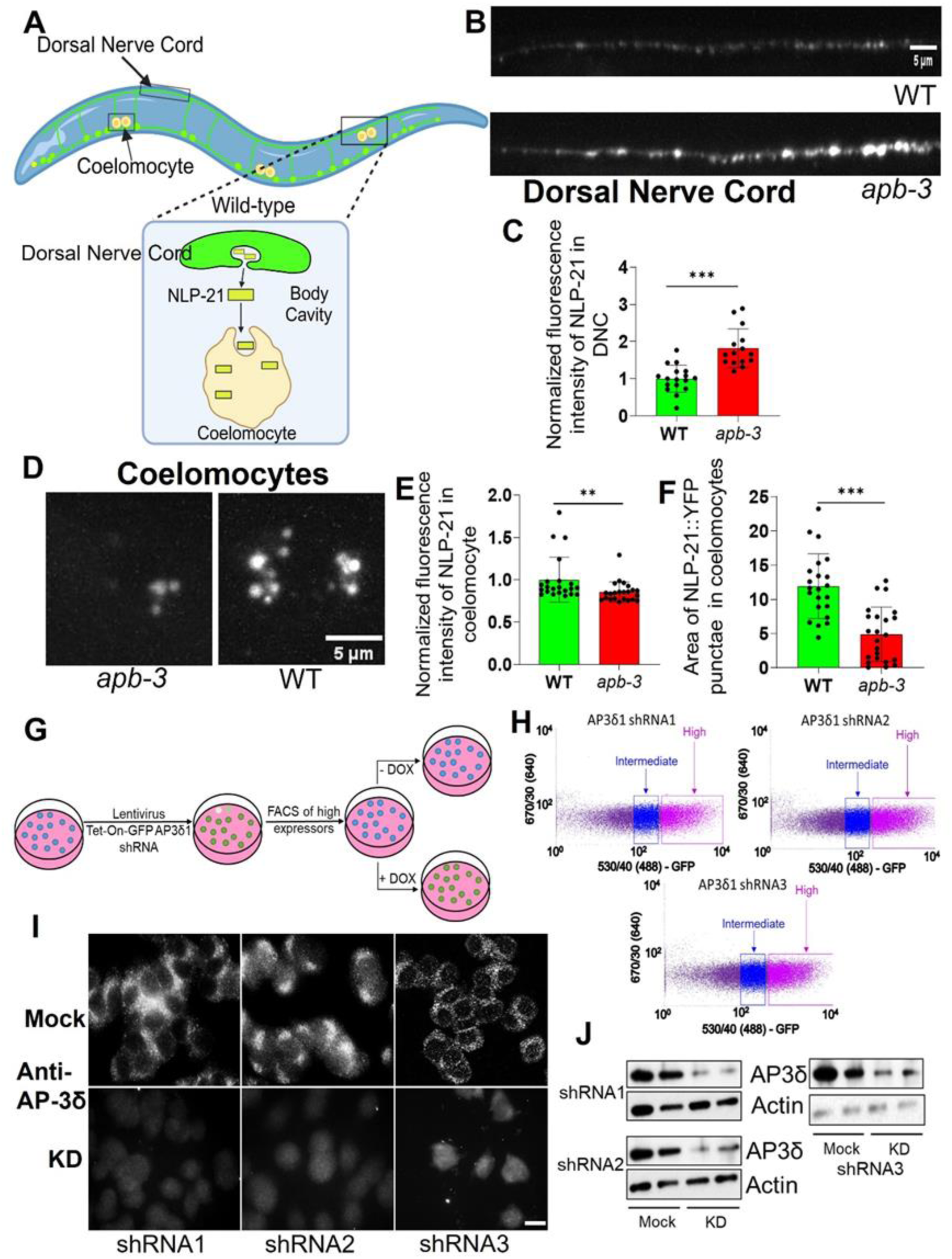
Impaired DCV exocytosis of NLP-21 in *C.elegans* and generation of inducible AP-3 knockdown in PC12 cells. (A) Schematic showing the secretion and endocytosis of NLP-21 from the posterior Dorsal Nerve Cord (DNC) and Coelomocytes, respectively. (B) Representative images showing NLP-21::YFP intensity near the posterior DNC in WT and apb-3(ok429) mutant animals. (C) Quantitation of mean intensity of NLP-21::YFP near the posterior DNC in WT (n=16) and apb-3(ok429) (n=15) mutant animals. Fluorescence intensity normalized to the wild-type for comparison (Mann– Whitney U test two-tailed). (D) Representative images showing NLP-21::YFP intensity in the posterior coelomocytes in WT and apb-3(ok429) mutant animals. (E) Quantitation of mean intensity of NLP-21::YFP above threshold in the posterior coelomocytes in WT (n=22) and apb-3(ok429) (n=23) mutant animals. Fluorescence intensity was normalized to the wild-type for comparison (Mann–Whitney U test two-tailed). (F) Quantitation of area covered by NLP-21::YFP punctate above threshold in the posterior coelomocytes in WT (n= 22) and apb-3(ok429) (n=23) mutant animals. Fluorescence intensity normalized to the wild-type for comparison (Unpaired Student’s t-test two-tailed). (G) Schematic of the protocol. Using the SMARTvector lentiviral system, cells were transduced with lentiviral shRNA (sh1, sh2 and sh3) targeting different sites for the Adapter Protein-3 Complex Delta1 subunit (AP3δ). As the shRNA is inducibly coexpressed with GFP upon treatment with doxycycline, cells were treated for 24 hours with doxycycline. After 24 hours, the doxycycline was removed, and the high-expressing cells were sorted using flow cytometry (H); selection was carried out with puromycin to obtain a stable population of cells with inducible AP3 shRNAs. To validate AP3 KD, cells were mock-treated or treated with doxycycline for 3 days. Cells subjected to Immunofluorescence (I) and Western blotting (J) show a robust loss of AP3 using antibody specific to the δ subunit of AP-3. Scale bar: 10 μm.

### Generation and validation of stable and inducible AP-3 knockdown PC12 cells

AP-3 is a heterotetrameric complex where targeting the δ subunit allows a true depletion of the AP-3 complex as the δ subunit trunk forms the core of the AP3 complex^22,37^. Based on the preliminary C.elegans data, we developed a doxycycline-inducible Tet-on knockdown system in PC12 cell lines against the δ subunit of AP-3 to enable the temporal loss of the AP3 complex upon a five-day treatment. We designed three unique guides targeting different regions of the AP-3 δ subunit, and a mixed population of cells with high expression were FACS sorted and stably selected (Fig 1G, H), established as knockdown cells for all subsequent experiments (See materials and methods). All three cell lines showed a significant depletion in the expression of AP3 (Fig 1I, J) upon induction with doxycycline treatment in immunofluorescent images and immunoblots, respectively.

### Stimulus-coupled secretion experiments and Biochemical analysis reveal compromised DCV exocytosis in AP3-depleted cells

In line with these findings, immunoblotting experiments documented a significant impediment in the release of CgB (chromogranin B), a DCV marker^38^ upon depolarization of AP-3 depleted PC12 cells using multiple shRNA when compared to the mock cells where the release of CgB is unaffected (Fig 2B – G, Supplementary Fig 1A-B). We also noted a marginal increase in the protein level of CgB (Fig 2B – G) but not at the transcript level, suggesting a disruption in cargo release from the DCVs concerning the Mock; a similar finding is reported^21^. These findings could be attributed to previous findings where the loss of AP-3 increases DCV size and volume^19^. To rule out any defect in constitutive release, we observed no changes in protein levels of HSP90 in the supernatant (specific marker) or global constitutive signature^39,40^ (Supplementary Fig 1C-G). We also used a scrambled shRNA PC12 cell line as a control, indicating no changes in the release of CgB (Supplementary Fig 1H). To substantiate our claim, we focused on the regulated exocytosis of the proteolytic processed fragment of Peptidylglycine alpha-amidating monooxygenase (PAM), an integral membrane enzyme resident on DCVs^35,31,40,41^. Mock cells, when subjected to secretagogue coupled secretion, show a distinct release of sPHM fragment typically processed in the DCVs in the supernatant, which was hugely diminished in the supernatant in AP-3 depleted cells (Fig 2J-K, Supplementary Fig 1K-L).

**Figure 2.**
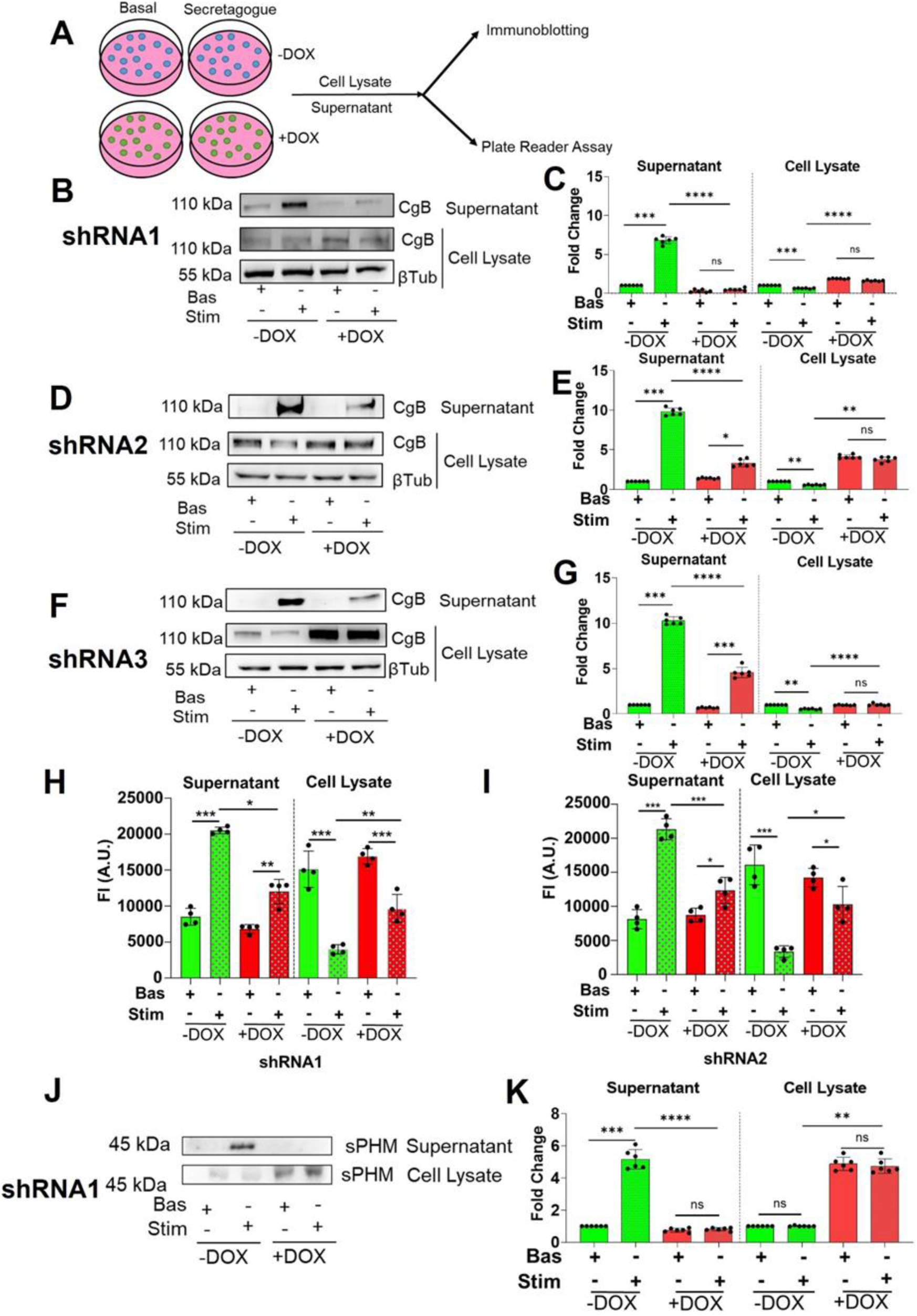
AP-3 knockdown in PC12 cell lines causes an impediment in the regulated exocytosis of dense core vesicles. (A) Schematic diagram of the protocol. Secretion of Chromogranin B (CgB) in mock-treated and AP3-depleted cells (treatment with doxycycline for 3 days). Western blot, quantification (B-G, for the three separate shRNA) for the mock-treated, and AP3 depleted cells incubated with BaCl2 for 15 mins, post that both the cells and supernatants from the cells were probed with anti-CgB. AP3 depletion impedes the secretion of CgB. (H, I) Fluorescent intensity was measured using a plate reader of mock-treated and AP3-depleted cells (shRNA1 and shRNA2) transfected with NPY-mApple, and n=3 showed depletion in NPY release. The release of sPHM from mock-treated (Green) and shRNA1 AP3 depleted cells (Red) upon stimulation with a secretagogue KCl, (J-K) WB, and quantification shows a major defect in release (n=3). Data is shown as mean ± SD. (*P<0.05, **P<0.01, ***P<0.001; n.s. statistically non-significant).

Interestingly, we could note an increased presence of sPHM in cell lysates even on secretagogue stimulation (Fig 2J-K, Supplementary Fig 1K-L), denoting a defect in the regulated release of PAM post cleavage. In parallel, NPY (Neuropeptide Y, a soluble DCV cargo) tagged with a fluorescently labelled mApple plasmid was introduced in AP3-depleted cells to generate stable cells. Stimulus-coupled secretion experiments were performed in Mock and AP3-depleted cells, followed by measuring fluorescent intensities in the supernatant and lysates (shRNA1 and shRNA2 cells) (Fig 2H-I). Results indicated defective regulated secretion, as evidenced by the reduced fluorescent NPY levels upon stimulation in supernatants of AP3-depleted cells. Meanwhile, in mock cells, fluorescence intensities of NPY increased in the supernatant, corresponding to an unaffected regulated release of DCVs (Fig 2H-I). Thus, in conjunction, these experiments suggest an impeded regulated secretory pathway due to the loss of AP-3.

### Live cell imaging suggests that DCV exocytosis is compromised

In mocha mice, the effects of lack of AP3 on DCV exocytosis are not well characterized^19,20^. To understand the dynamics of DCV exocytosis, we looked at individual exocytotic events in “Mock” and “AP3KD” conditions using NPY - pHTomato^30^ (a bonafide DCV cargo with a PH-sensitive fluorescently tagged RFP). KCl, being a secretagogue, induces membrane depolarization, which causes fusion and subsequent exocytosis of DCVs (Fig 3A). Upon stimulation, we observed an increased NPY-pHTomato fluorescence in mock-treated cells, while shRNA1 AP3-depleted cells showed no observable response (Fig 3B-C, S1, S2). We also observed no significant change in the basal fluorescence of NPY-pHTomato among WT and shRNA1 AP3-depleted cells (Fig 3D), suggesting no defective synthesis of the Cargo. As a measure of DCV exocytosis, we quantified the maximum change in NPY-pHTomato puncta fluorescence (maximum Δf/f0), which shows a significantly diminished release of overall vesicular content in AP3-depleted cells (Fig 3E). The responsiveness of vesicles consisting of NPY-pHTomato was assessed by counting individual exocytotic events^39^ with a minimum Δf/f0 of 0.3 as the threshold. From Fig (3F-G), an overall decline in responding vesicles further indicates that the exocytosis of DCVs is heavily impaired in case of AP3 depletion. We also observed similar impairment in real-time exocytosis of DCVs in shRNA2 AP3 depleted cells compared to mock-treated cells (Supplementary Fig 1M-1O, S3, S4), ruling out any off-target effects and reconfirming the robust phenotype. Hence, our imaging experiment is more valuable in determining a defective DCV exocytosis.

**Figure 3.**
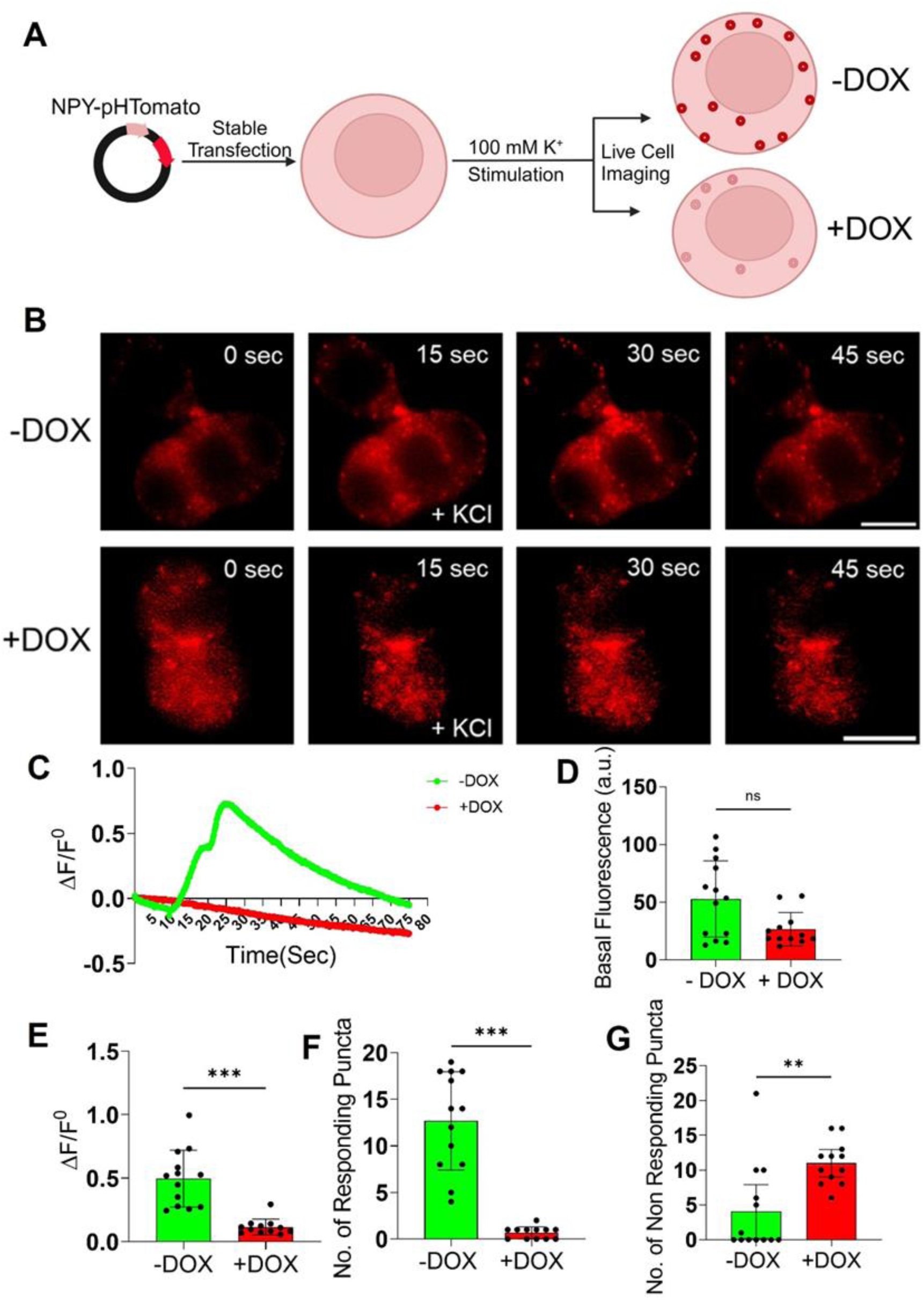
AP-3 knockdown causes an impairment in the real-time exocytotic events in DCVs. (A) Schematic showing stable transfection of NPY-pHtomato and the protocol for live-cell imaging experiment. (B) Representative images of mock-treated and shRNA1 AP3 depleted cells transfected with NPY-pHtomato (red) showing basal and stimulated conditions. Scale bar 5 µm (C) Representative time traces of mean NPY-pHtomato fluorescence intensities showing an average of all puncta per cell (green, mock-treated; red, shRNA1 AP3 depleted cells). (D) Represents no change in the basal fluorescence of NPY-pHtomato in mock vs AP3 depleted cells. (E) Quantitative analysis of the maximum Δf/f0 in mock-treated (n=13 cells) and shRNA1 AP3 depleted cells (n=12 cells). (Mann-Whitney Test). (F) Number of responding NPY-pHtomato puncta/cell (Mann-Whitney test) (G) Number of non-responding NPY-pHtomato puncta/cell (Mann-Whitney test) (F, G) Considering a minimum of Δf/f0 of 0.3 upon KCl stimulation as a response. (N=4). (‘N’ denotes the number of independent experiments, and ‘n’ is the number of cells taken for quantification) Data are shown as mean ± SD. (*P<0.05, **P<0.01, ***P<0.001; n.s. statistically non-significant).

### Perturbed release of SLV cargo Synaptophysin upon stimulation

PC12 cells harbour vesicles identical to synaptic vesicles in neurons originating from early endosomes and are referred to as synaptic-like vesicles (SLVs)^41^. Given the heavy localization of AP3 on endosomes and previous reports pointing towards the requirement of neuronal AP3 for SLV formation^9^, we tested the functional status of SLVs using pHTomato^42,43^ (Fig 4A). Synaptic vesicle exocytosis was normal in WT-type cells with a mild perturbation in AP3-depleted cells (Fig 4B-C, S5, S6). We observed no significant change in the basal fluorescence of Synaptophysin-pHTomato among WT and shRNA1 AP3-depleted cells (Fig 4D). The maximum change in fluorescence (maximum Δf/f0) for Synaptophysin-pHTomato puncta, similar to DCV exocytosis, was used to quantify SLV exocytosis. A significant reduction in maximum Δf/f0 indicates a defective SLV exocytosis in AP3-depleted cells (Fig 4E). The increase in non-responsive Synaptophysin-pHTomato puncta and the corresponding decline in responding puncta substantiate the SLV exocytosis defect (Fig 4F, G). These findings are of interest and are in line with previous findings^44^. However, SLVs significantly differ from synaptic vesicles in terms of compartmentalization and function.

**Figure 4.**
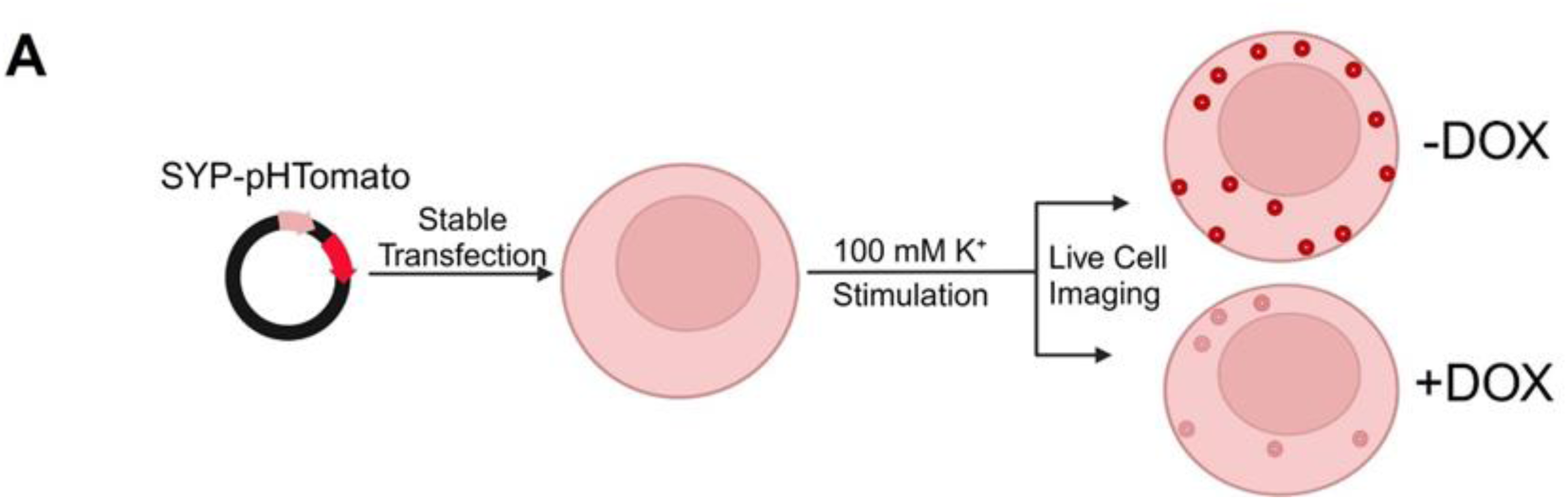
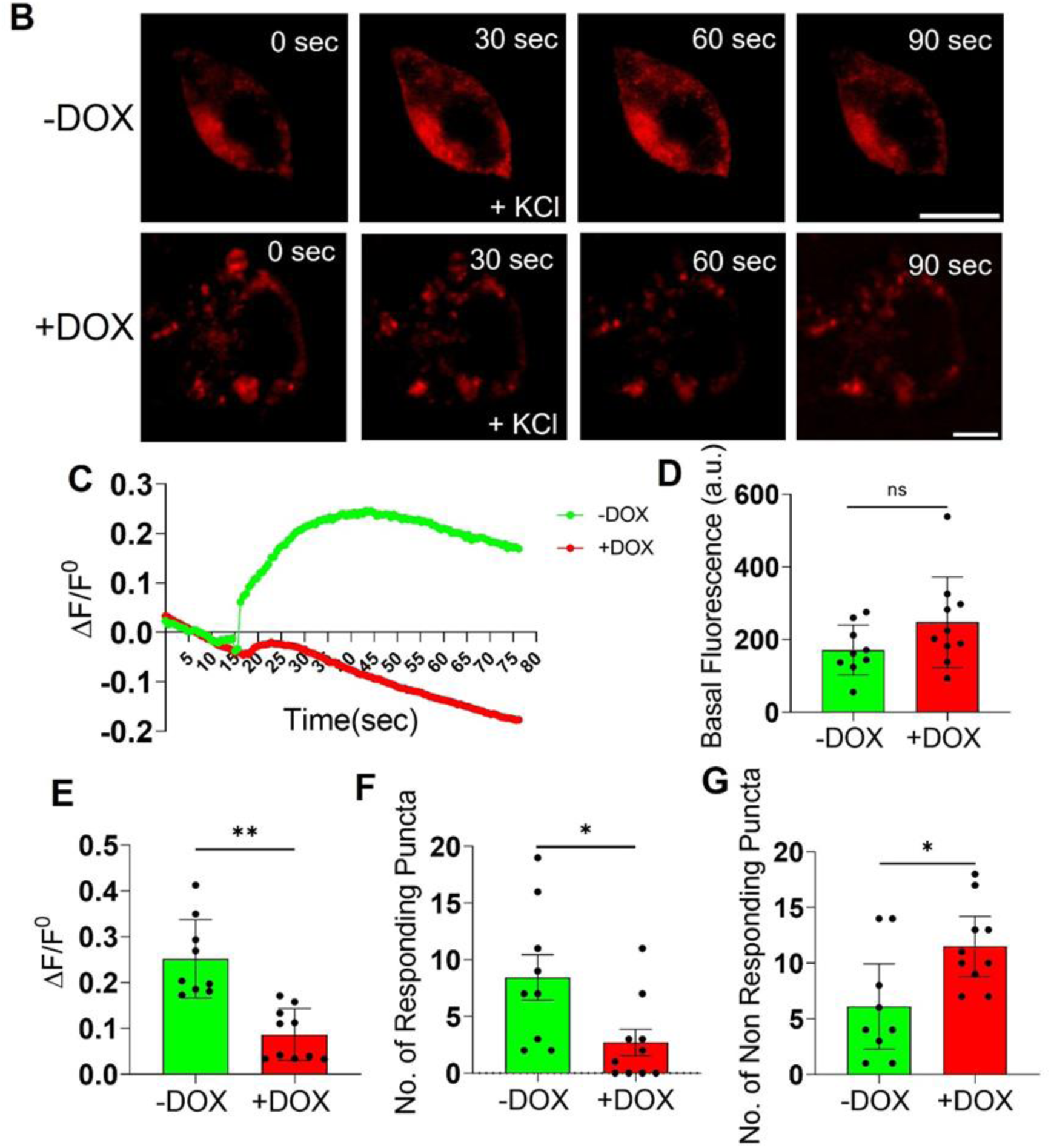
AP3 depletion causes an impairment in the real-time exocytotic events in Synaptic vesicles. (A) Schematic showing stable transfection of Synaptophysin-pHtomato and the protocol for live-cell imaging experiment. (B) Representative images of mock-treated and shRNA1 AP3 depleted cells transfected with Synaptophysin-pHtomato (red) showing basal and stimulated conditions. Scale Bar 5µm (C) Representative time traces of mean Synaptophysin-pHtomato fluorescence intensities showing an average of all puncta per cell (green, mock-treated; red, shRNA1 AP3 depleted cells (D) Represents no change in the basal fluorescence of Synaptophysin-pHtomato in Mock vs AP3 depleted cells. (E) Quantitative analysis of the maximum Δf/f0 in mock-treated (n=8 cells) and shRNA1 AP3 depleted cells (n=8 cells). (Mann-Whitney Test). (F) Number of responding Synaptophysin-pHtomato puncta/cell (Mann-Whitney test) (G) Number of non-responding Synaptophysin-pHtomato puncta/cell (Mann-Whitney test) (F-G) considering a minimum of Δf/f0 of 0.2 upon KCl stimulation as a response. (N=3). (‘N’ denotes the number of independent experiments, and ‘n’ is the number of cells taken for quantification) Data are shown as mean ± SD. (*P<0.05, **P<0.01, ***P<0.001; n.s. statistically non-significant).

### Ultra-structural analysis reveals that the maturation of DCVs is compromised

A previous report shows that decreased exocytosis can indicate possible maturation defects^24^. In conjunction with a previous Electron microscopy report^19^, We see a bigger dense core (Fig 5 (A-B) and Supplementary Fig 2 (C-D) while the overall size of DCV remained unchanged(Fig 5C and Supplementary Fig 2E). Furthermore, there was a decline in the DCV numbers (Fig 5D-E and Supplementary Fig 2F) in AP3 depleted cells. Images revealed that DCVs are distantly located from the plasma membrane in AP3KD cells (Fig 5F, G and Supplementary Fig 2G, H) while most of these vesicles are present closer to the Plasma membrane in WT conditions. In keeping with conventional views^19,45^, Increased volume indicates immature vesicle formation due to a lack of functional AP3 complex and accumulation of such vesicles at the proximity of TGN, resulting in an overall reduction of readily releasable pools of DCVs.

**Figure 5.**
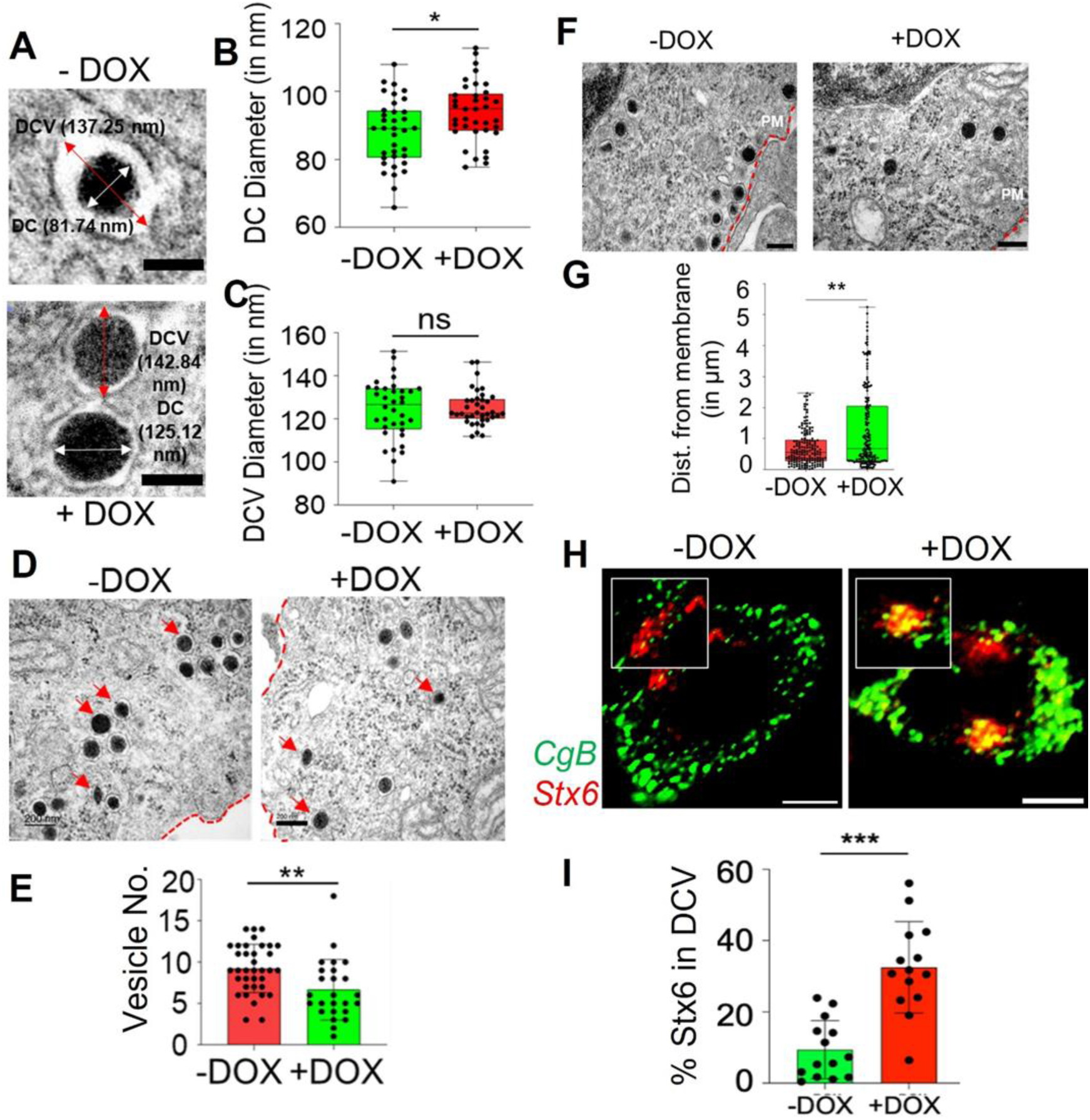
AP-3 affects the maturation of DCVs in PC12 cells. Electron microscopy of mock-treated (Green) and shRNA1 AP-3 Depleted cells (Red). (A-C) Representative images of the Dense core vesicles and the graphs show an increase in DC diameter but an unaffected DCV diameter. Scale bar: 100nm. (D-E) Representative images for the number of vesicles and a graph indicating a decline in the overall number of vesicles. Scale bar: 200nm. (F-G) Representative images for the DCV distance from the membrane and the graph show that the vesicles are farther away from the membrane in the case of AP3 KD. Scale bar: 200nm. (H-I) Representative confocal images and quantification showing an increase in colocalization (% colocalization) of Stx6 (Red) with CgB (DCV marker) (N=3, n=14 cells each) (Green). (n=9-10 cells for each condition, *P<0.05, **P<0.01, ***P<0.001; n.s. statistically non-significant).

### Enhanced localization of Stx6 to DCVs substantiates maturation defects

A previous study found that Syntaxin 6 (Stx6) is present in immature DCVs (dense core vesicles); it was further demonstrated that clathrin/AP-1 helps remove Stx6 from these vesicles and return them to the TGN (trans-Golgi network). This suggests that mature DCVs have very little or no Stx6^46^, and Stx6 is a signature protein of DCVs. Therefore, we wanted to explore if Stx6 would be localized more to the DCVs without AP-3. Hence, we compared the colocalization of Stx6 with chromogranin B (DCV marker) in mock and KD cells. Loss of AP-3 led to a two-fold increase in the colocalization of Stx6 with CgB (Fig 5H-I and Supplementary Fig. 2I-J), suggesting a defect in the removal of Stx6 from the immature DCVs that we see accumulated Stx-6 in AP-3 depleted cells. Given the well-known redundant functions of AP complexes^47^, AP-3 may also have a role in removing Stx6 from these vesicles and shuttling back to TGN, thus playing an additional role in DCV maturation by ‘sorting out’ the Cargo away from DCVs similar to the known role of AP-1 complex^46^.

### RUSH (Retention under selective hook) experiments reveal slower Kinetics of DCV budding at Golgi in AP3-depleted cells

Beyond its contribution to DCV maturation, we sought to explore the broader spectrum of AP-3 functions at the TGN. The journey towards becoming mature DCVs starts post-TGN budding.^48^. AP-3 dependent budding of DCVs and Cargo sorting at the TGN is poorly understood. Hence, we used the Retention Using the Selective Hook^49^ approach to trace the trafficking of a soluble DCV cargo, Neuropeptide Y (NPY), from ER through Golgi to the DCVs. To begin with, NPY is present throughout the cytoplasm without biotin, consistently localizing with the ER. Treatment with biotin releases the NPY from the ER hook, leading it towards the Golgi, where it localizes completely with a TGN marker Golga 1(Golgin 97) at 20 mins post-treatment in both Mock and AP3 depleted PC12 cells (Fig 6B-D). While the budding of NPY-bearing vesicles from the TGN was majorly accomplished (approximately 70%) by 40 mins, we see complete NPY exit from the TGN by 60 mins in mock cells (Fig 6B and 6D). This NPY-RUSH phenomenon has been established in the case of PC12^42^, which concurs with our data in PC12 cells. On the contrary, we observed slow budding of NPY-bearing immature DCVs as by 40 mins, NPY had no change in its localization with TGN. However, at 60 minutes, a clear redistribution of NPY in the cytoplasm as the colocalization with TGN starts to decrease (Fig 6C-D) in AP-3 depleted cells. We have also estimated the rates at which NPY enters and exits the Golgi (described in methods) and observed no changes in the rate of entry of NPY into the Golgi (Fig 6E). However, we see a significant decline in the Rate of exit of NPY from the TGN in the depletion of AP-3 (Fig 6F). Similar observations were documented in the case of AP3δ shRNA2 harboring cells (Supplementary Fig 3A-E), strengthening the role of AP3 in the budding of DCVs at the TGN and a delayed budding of these vesicles in the absence of AP-3.

**Figure 6.**
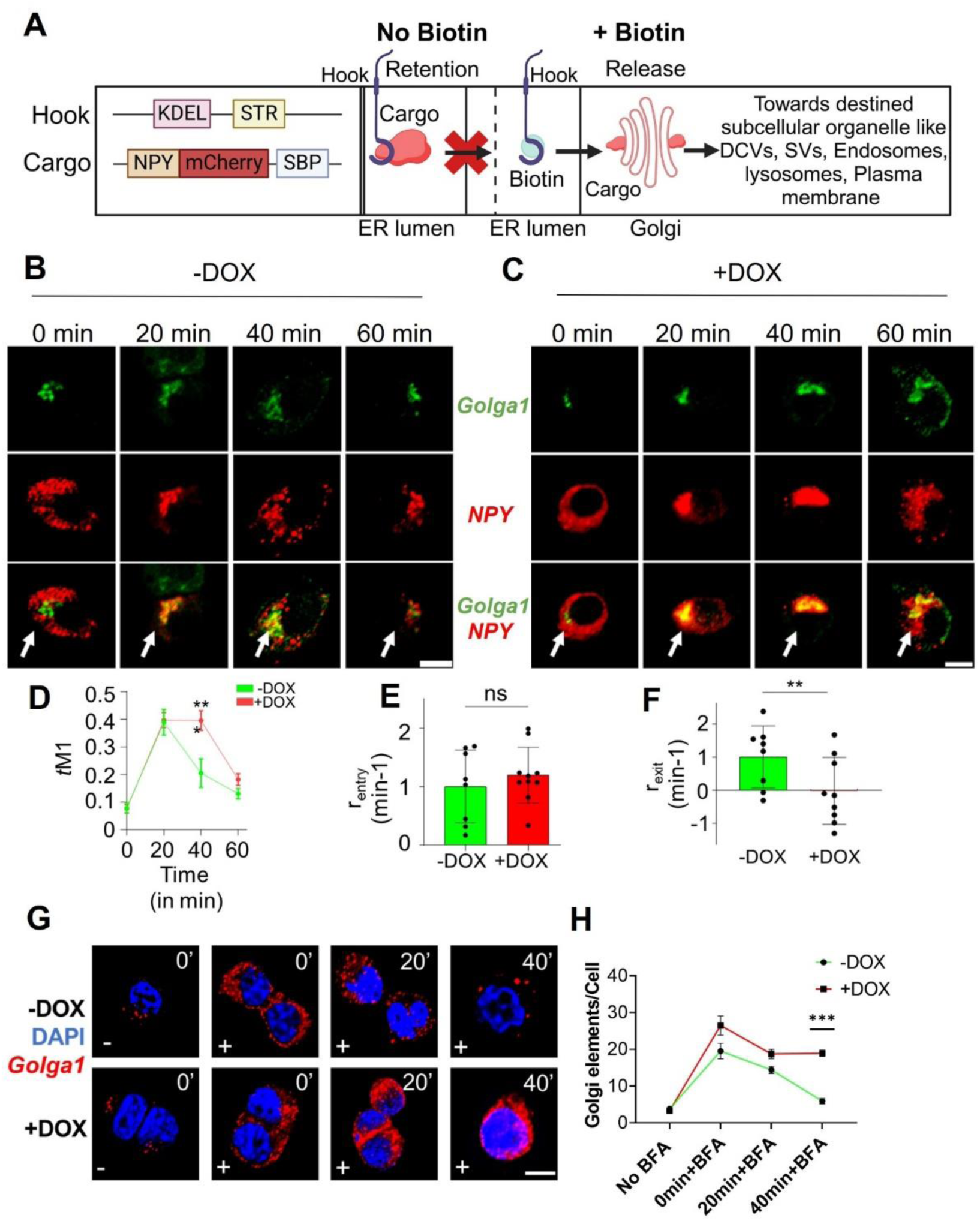
AP3 affects the dynamics organization of the trans-Golgi network and its subsequent sorting of DCV cargoes. (A) Schematic of Retention Using Selective Hook (RUSH) system. (B-C) Representative confocal microscope fluorescent images for mock-treated and shRNA1 AP3 depleted cells, where Golga1 is green and NPY is red. Scale bar: 5µm (D) Graphs showing tM1 (Mander’s coefficient 1) of NPY-mCherry and Golga1 for mock-treated (Green) and shRNA1 AP3 depleted cells (Red). (E) Graphs for the Rate of entry of Cargo at the Golgi per minute. (F) Graphs show a reduction in the Rate of cargo exit from the Golgi per minute due to AP3 KD. (N=3 independent experiments, n=10 cells) Scale bar: 5µm. (G-H) Representative images and quantification for mock-treated and shRNA1 AP3 depleted cells treated with Brefeldin A showing a defect in the Golgi reassembly upon AP3 KD after 40 minutes of Bfrefeldin-A washout. (Each data point represents a different field of cells containing 6-31 cells per field, where each condition has 9 fields)(N=3) Scale bar: 5µm. (*P<0.05, **P<0.01, ***P<0.001; n.s. statistically non-significant).

### Brefeldin A experiments suggested dysfunctional Golgi

AP-3 is a coat protein dependent on ARF1 and Arf-GAP AGAP1 for its binding with Endolysosomal membranes, where ARF1 inhibited by Brefeldin A (BFA) leads to Golgi disassembly^13,50^. Therefore, we wanted to visualize the dynamic organization of the Golgi and specify any involvement of AP-3 in the Golgi function. Brief incubation with BFA for 15 mins shows a disruption of Golgi that reversibly begins to reassemble after washout of BFA in mock PC12 cells based on the staining with the TGN marker, Golgin 97. At 40 minutes post washout of BFA in mock cells, TGN elements per cell declined, suggesting an overall reassembly of the organelle (Fig 6G-H and Supplementary Fig 3F).

Interestingly, in AP-3 depleted cells, Golgi disassembly was irreversible, i.e. post wash out, no significant reduction in Golgi fragmentation was observed (Fig 6G-H and Supplementary Fig 3F. It had been previously reported that a similar defect in the reassembly of Golgi was observed in the case of loss of clathrin heavy chain ^51^. This may suggest that AP-3, along with the clathrin heavy chain, also plays an important role in the reorganization of the Golgi, and dysfunction can lead to disrupted sorting sites and cause missorting of Cargo being loaded onto the DCVs, which are being potentially retrieved by recycling.^52^

### Specific DCV cargo is depleted in the vesicle-enriched fractions of AP3 perturbed cells

AP-3 has been implicated in the sorting of various cargo proteins, such as Zinc Transporter-3 (ZnT-3) and chloride channel (ClC-3), which have been reported as signature proteins of both DCVs and SLVs (^53,54^). However, it is not well understood if AP-3 also sorts distinct or novel Cargo specific to each compartment. For this purpose, we used the Comparative proteomics approach to identify the various Cargo and other accessory proteins associated with AP-3. Mock and AP3-depleted cells were grown in “light” and “heavy” stable isotopes of amino acids (Arginine and Lysine) for three weeks. Post labeling, either cell lysates were fractionated by differential ultracentrifugation to obtain a vesicle-enriched fraction comprising mainly clathrin-coated vesicles, dense core vesicles, and Synaptic-like vesicles (Fig 7A, See methods). This method and the analysis were based on the previous works of ^24,55^. From the mass spectrometric analysis, we detected approximately 2000 proteins, including soluble and membrane proteins of DCVs represented as Scatter plots against -log2[H/L] values showing deficient proteins in VEFs (coloured dots represent significantly depleted proteins) for Mock vs shRNA1 AP3 depleted cells (Fig 7B). Clathrin heavy chain (CHC) is the most abundant protein in the vesicle-enriched fraction and serves as a control for this fractionation experiment^55^. CHC was enriched in the fractions with unaltered Mock and AP3 depleted conditions. AP-3 is a coat protein complex functioning both in a clathrin-dependent and independent manner, explaining the unaffected levels of clathrin due to this partial dependence^56,57^ (Fig 7C & Supplementary 4A). 66 proteins were significantly reduced in AP-3 depleted cells compared to mock cells in the vesicle-enriched fractions (Supplementary Table 1).

**Figure 7.**
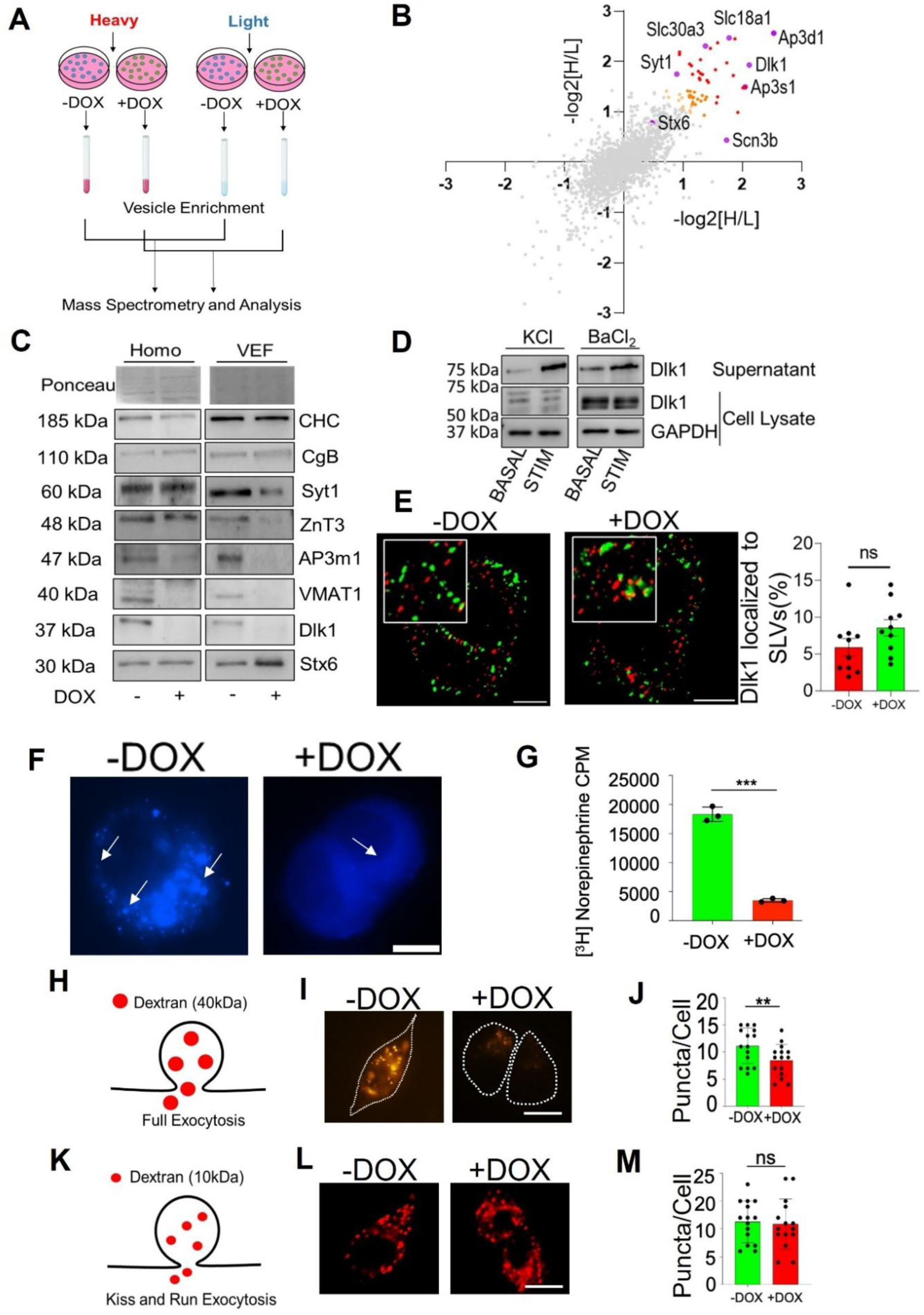
AP3 plays a selective role in sorting membrane proteins and also affects the trafficking of vesicles post-Golgi. (A) Schematic of the SILAC-based Mass Spectrometry approach to assessing VEFs for mock-treated and shRNA1 AP3 depleted cells. (B) Scatter plots against -log2[H/L] in 2 separate experiments show deficient proteins in VEFs (coloured dots represent significantly depleted proteins) for Mock vs shRNA1 AP3 depleted cells. These deficient proteins include certain membrane proteins known as DCV cargoes or novel cargo proteins for AP3-dependent vesicles. (C) WB validates the abundance in the Cell homogenate and VEFs for the proteins deficient in MS in either condition. (D) Western blot for the WT PC12 cells incubated with BaCl2 for 15 mins and KCL for 20 mins, post that both the cells and supernatants from the cells were probed with anti-Dlk1. (E) Representative confocal images showing colocalization of Dlk1 (Red) with synaptic vesicle marker synapsin-2 (Green) in Mock vs shRNA1 AP3 depleted cells showed no significant colocalization change. Scale bar 5um (n=10 cells each) (F) Representative fluorescent images of the mock-treated and shRNA1 AP3 depleted cells showed reduced Zinquin ethyl ester dye uptake. Scale bar: 5µm. (G) The graph depicts approximately five-fold more uptake of Nor-epinephrine in the mock-treated cells after adding up the counts in both cells and the medium. (***Indicates statistical significance at the *p* < 0.001 level assessed by two-tailed Student’s *t*-test (three independent experiments). (H) Schematic for the full exocytotic events. (I-J) Representative images and quantification for Dextran 40kDa show impaired full fusion exocytosis as the puncta per cell reduces in shRNA1 AP3 depleted cells (Red) w.r.t Mock treated cells (Green). (N=3, n=15 cells each). (K) Schematic for the Kiss and Run Exocytotic Events. (L-M) Representative images and quantification for Dextran 10kDa show no significant changes in mock (Green) vs shRNA1 AP3 depleted cells (Red). (N=3. N=15 cells each). *P<0.05, **P<0.01, ***P<0.001; n.s. statistically non-significant).

Some of these candidates for which antibodies were available were validated by western blotting (Fig 7C & Supplementary Fig 4A), indicating clear enrichment of Stx-6 in the vesicular fraction that strengthens our claim in suggesting AP-3’s involvement in transporting Stx6 back to the TGN from the immature DCVs. Other AP-3 complex subunits, such as AP3 β and σ, have also been reported to be depleted, while the µ subunit was detected only in one run. Based on our data from (Fig 2 B, D, F), we showed an increase in the CgB levels in the whole cell lysate, with a 32% increase in the vesicular fractions in the absence of AP-3 (Fig 7C). This infers an accumulation of cargoes in the immature DCVs, causing inefficient exocytosis.

Among the significantly depleted proteins, we report the presence of multiple ion channels (Vdac1, Scn3b) and transmembrane proteins (Dlk1, VMAT1, ZnT3, Syt-1), which could be dependent on AP-3 complex for their sorting. Specifically, we highlight Synaptotagmin-1(Syt1), which functions as a calcium sensor helping in vesicle fusion and neurotransmitter release from DCVs^25^ reduced by 79% in the AP3 depleted vesicle fraction (Fig 7C & Supplementary Fig 4A). At the same time, its levels in cell homogenate remained unchanged, a finding consistent with previous reports ^21^. Similarly, Vesicular monoamine transporter 1 (VMAT1), another DCV cargo responsible for the loading of catecholamines into the DCVs^26^ is almost entirely lost from the vesicular fraction as well as whole cell homogenate in the absence of AP-3 (Fig 7C & Supplementary Fig 4A). This suggests a possible involvement of AP-3 in sorting VMAT1 towards the regulated secretory pathway. We also report reduced abundance levels of ZnT3 in the vesicular fractions by 88% (Fig 7C & Supplementary Fig 4A), a finding consistent with previous reports^53^.

Interestingly, we report that Delta-like homologue1 (Dlk1), a non-canonical notch ligand-receptor, also disappears from vesicular and whole cell fractions (Fig 7C). It is noteworthy from a previous report that the absence of Dlk1 transcript in a clonal PC12 cell line has defective regulated secretion ^58^. Here, we are reporting Dlk1 as novel DCV cargo, which is secreted in a regulated manner based on the secretagogue-based release of Dlk1 upon stimulation (Fig 7D). As we demonstrated a clear loss of Dlk1 in the absence of AP-3, we suggest Dlk1 is a possible novel AP-3 dependent DCV cargo. Likewise, we show that Dlk1 is not present on the synaptic vesicles (Fig 7E), implying that Dlk1 is specific to DCVs.

### Mislocalization of ZnT3 and VMAT1 corroborates with functional defects in the accumulation of neurotransmitters and Zinc in the DCV compartment

Building on the proteomics data, we looked at the functions associated with the proteins depleted in vesicle-enriched fractions. The function associated with ZnT3 is the accumulation of Zinc inside SLVs and DCVs^59–61^. Therefore, we used Zinquin ethyl ester (ZEE)^62^, a dye that specifically binds to the zinc ions inside the vesicles. Mock cells incubated with zinc ions when stained with ZEE show distinct punctate staining, signifying Zinc uptake and mobilization, whereas AP-3 depleted cells did not show any puncta, indicating defective uptake of Zinc (Fig 7F and Supplementary Fig 4B). The disappearance of ZnT3 due to AP-3 depletion possibly accounts for this defect. The uptake and release of catecholamines (nor-epinephrine, epinephrine, dopamine) depends on a transporter VMAT1 found primarily on regulated secretory vesicles. Therefore, we investigated nor-epinephrine secretion using a catecholamine uptake assay.^24^ (see methods). This showed us that loss of AP-3 led to a five-fold reduction in nor-epinephrine release compared to mock cells (Fig 7G and Supplementary Fig 4C). This again corroborates nicely with reduced VMAT-1 from the AP-3 depleted vesicle enriched fractions. Hence, when AP-3 is depleted from these cells, the localization of ZnT3 and VMAT1 may be lost, signifying a probable sorting defect. AP-3 is also localized at the early and recycling endosomes, regulating the trafficking of the cargoes to the lysosomes or lysosome-related organelles^63^. Hence, we validated the defective Endolysosomal functioning in case of AP-3 depletion using the dextran 10kDa (experiment described in methods), implying a delay in the clearance after 30 mins and 60 mins of dextran uptake (Supplementary Fig 4G, 4H). Thus, this signifies the importance of AP-3 in regulating exocytotic recycling and clearance via the Endolysosomal pathway.

### Depletion of Syt1, a key protein of the SNARE complex, corresponds to defective full fusion exocytosis

There are multiple modes of regulated exocytosis: kiss-and-run exocytosis, which involves transient fusion of the secretory granule with the plasma membrane followed by its recycling; and full fusion exocytosis, which involves full fusion of a single secretory granule with the plasma membrane.^64^. Smaller pores around 5 nm are involved in ’kiss and run’ events, allowing for the uptake of 10kDa Dextran molecules. Conversely, larger pores, formed during full fusion events and measuring 9 nm in diameter, are responsible for the uptake of 40kDa Dextran molecules^65–67^. Mock and AP-3 depleted cells were exposed to two different sizes of rhodamine-Dextran, 10 kDa and 40 kDa, for one minute in an environment with high potassium concentration, allowing the dextran molecules to percolate into the cells through two mechanisms: kiss-and-run and full fusion, respectively. Loss of AP-3 led to a decline in the number of puncta with 40 kDa Dextran, signifying an impaired full fusion exocytosis (Fig 7H-J & Supplementary Fig 4C-E). Therefore, depletion of Synaptotagmin-1 (Syt1) justifies this impairment as Syt1 functions as a prominent Ca²+ sensor that critically regulates the final stage of vesicle fusion with the plasma membrane, facilitating full fusion events^68^. Meanwhile, the kiss and run exocytotic events were unaffected in AP-3-deprived cells (Fig 7K-M).

### Reduced localisation of Syt1, VMAT1 and DLK-1 with DCV compartment

We have shown that the lack of the AP-3 protein disrupts the transport of secretory vesicles besides making them non-functional. This is substantiated by the loss of key signature proteins such as Syt-1, VMAT-1, and ZNT-3, which are known to make the vesicles functional in various ways. We hypothesized that these proteins are perhaps being missorted, and to test this, we employed confocal microscopy to assess the colocalization of such proteins, which were significantly decreased and for which reagents were available. For instance, we looked at Syt1, Dlk1, and VMAT1 localization with the DCV marker CgB in mock and AP-3-depleted cells. Confocal microscopy revealed a near-complete loss of colocalization between Syt1, Dlk1, and VMAT1 with CgB^38^ in AP-3 knockdown cells (Fig 8A(A-F) & Supplementary Fig 5B(A-F)). This suggests that AP-3 is essential for sorting these known signature proteins into dense-core vesicles. To comprehensively evaluate the impact of AP-3 depletion on DCV cargo localization, we further investigated proteins identified by mass spectrometry as unaffected by AP-3 knockdown. We assessed the colocalization of neuropeptide Y (NPY) soluble cargo, phogrin, Synaptotagmin 4 (Syt4), Synaptotagmin 7 (Syt7), and vesicular monoamine transporter 2 (VMAT2) with the dense-core vesicle marker CgB in both mock-treated and shRNA1-mediated AP-3 depleted cells. Our findings revealed no significant differences in the colocalization patterns of these signature proteins amongst control and KD (Supplementary 5A), supporting our hypothesis that AP-3’s role in sorting specific cargo molecules destined for dense-core vesicles is preferential.

**Figure 8a.**
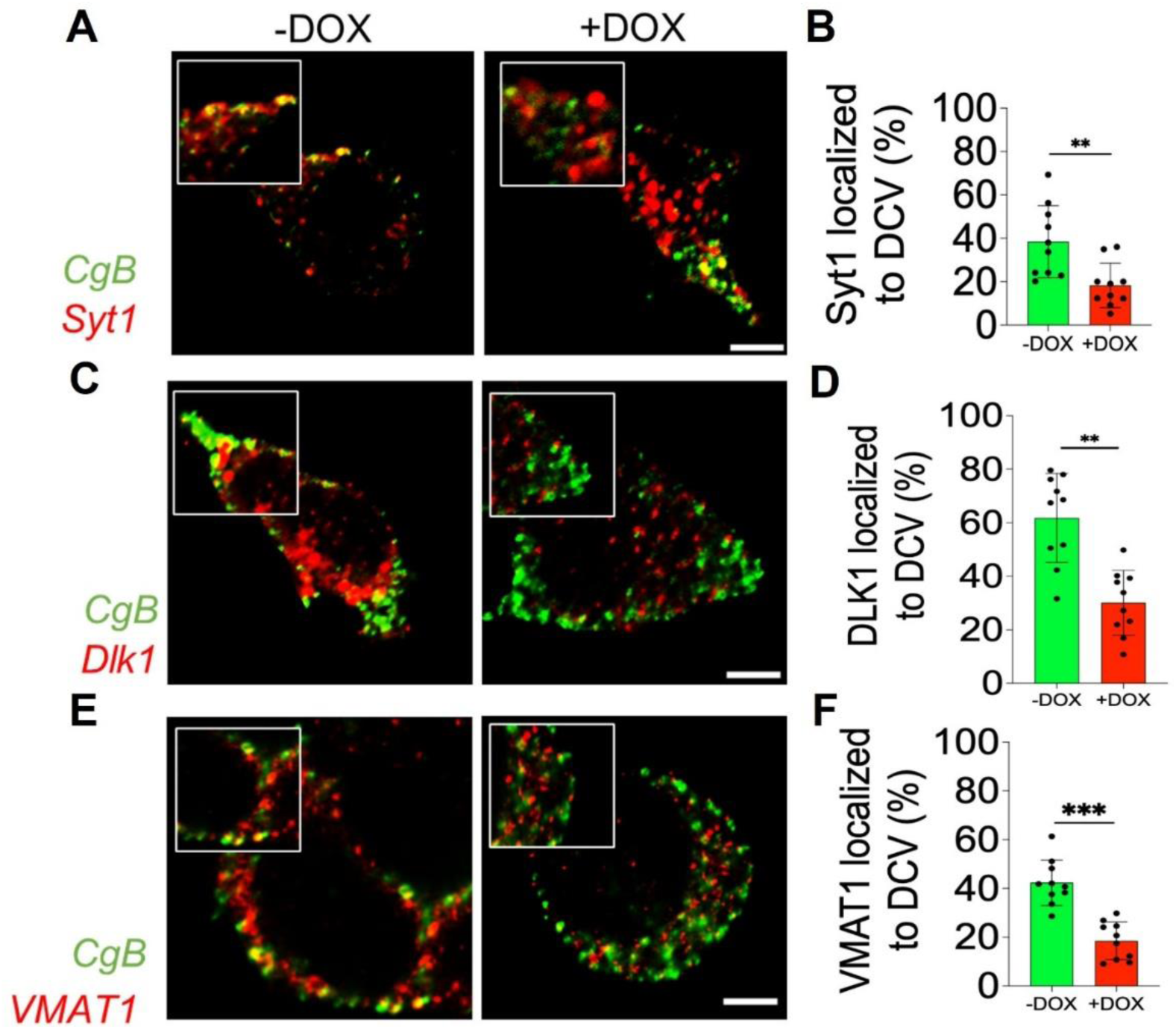
AP3 preferentially sorts specific membrane proteins mislocalized during loss of AP3 in PC12 cells. (A) Representative confocal images showing colocalization of specific proteins like Syt1, Dlk1 and VMAT1 with DCV marker CgB in Mock vs shRNA1 AP3 depleted cells. (A-F) Colocalization of Syt1, Dlk1 and VMAT1 (Red) with DCV(Green) reduces in AP3 knockdown. Scale bar 5um (n=10 cells or 3 independent experiments for both mock-treated and shRNA1 AP3 depleted cells, *P<0.05, **P<0.01, ***P<0.001; n.s. statistically non-significant).

### Augmented localisation of Syt1, VMAT1 and DLK-1 to lysosomes

Lysosomes act as the cell’s main recycling center, breaking down proteins that have reached their lifespan or are improperly folded or delivered within the cell. To further determine the fate of mislocalized proteins due to AP3 loss of function, we investigated the final destination of these cargo proteins by co-localizing them with a lysosomal marker (lyso 20 or LAMP1) in both control and AP3-depleted cells. AP3 knockdown significantly increased the colocalization of Syt1, Dlk1, and VMAT1 with lysosomal markers (Fig 8b(C-H) & Supplementary Fig 5c(C-H)). This approach revealed a potential misrouting of the cargo proteins towards lysosomes upon AP3 knockdown. Unlike Syt1, Dlk1, and VMAT1, the localization of CgB (a marker for DCVs) remained unchanged relative to lysosomes in AP-3 deficient cells (Fig 8b(A-B) & Supplementary Fig 5c(A-B)). This suggests that AP-3 deficiency does not affect the sorting of soluble Cargo such as CgB. Our findings suggest that AP-3 is critical in targeting specific membrane proteins (Syt1, Dlk1, and VMAT1) to DCVs. This mis-sorting possibly explains the observed depletion of these proteins upon AP-3 depletion.

**Figure 8b.**
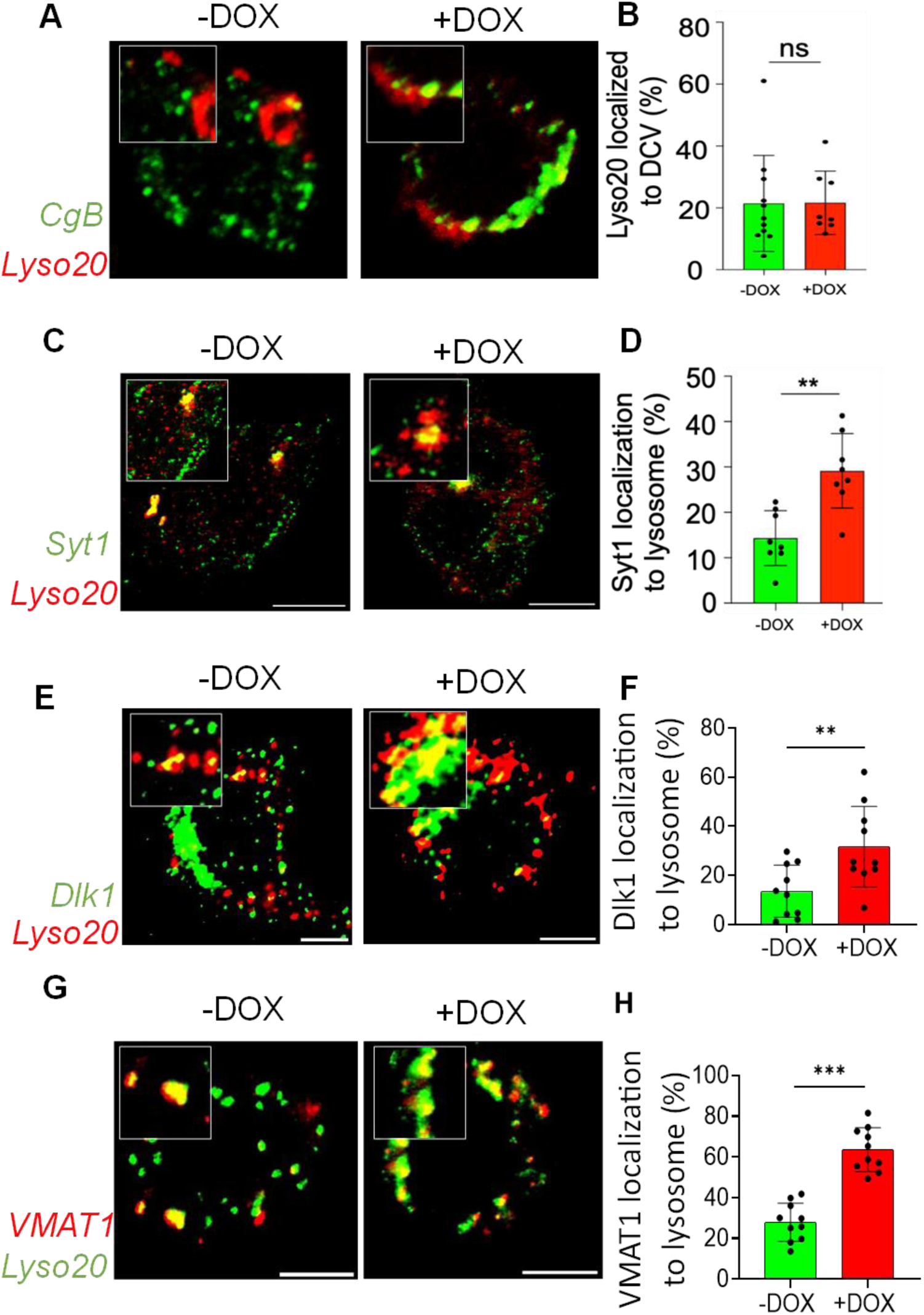
Missorted proteins are rerouted to lysosomes: Representative confocal images showing colocalization of Syt1, Dlk1, VMAT1, and CgB with Lysosomal marker LAMP1 in mock vs shRNA1 AP3 depleted cells. (A-B) Colocalization of DCV marker CgB (Green) with Lysosomal marker Lyso20 or LAMP1 (Red) shows no significant change. (C-J) Colocalization of Syt1, Dlk1 and VMAT1 (Green) with lysosomal marker Lyso20 or LAMP1 (Red) increases AP3 knockdown, indicating mis-sorting to the lysosomes. Scale bar 5um (n=10 cells or 3 independent experiments for both mock-treated and shRNA1 AP3 depleted cells, *P<0.05, **P<0.01, ***P<0.001; n.s. statistically non-significant).

### Computational analysis using Colabfold2 predicts physical associations of AP-3 with Syt1 and VMAT1

Loss of key specific DCV signature proteins in the vesicle-enriched fractions in AP3 depleted, accompanied by missorting and rerouting of these proteins to lysosomes, established the role of the AP 3 complex in sorting them to DCVs. We were curious to investigate if there are any physical interactions amongst these proteins with the AP 3 complex. This prompted us to use computational biophysics and a bioinformatics approach to predict the interactions amongst these proteins using a recently developed platform called ColabFold.^69^ We looked at candidate proteins that had been missorted in microscopy experiments and lost in vesicle-enriched fractions. The alphafold modelling compared all subunits of the AP-3 complex with Syt1, VMAT1, and Dlk1 protein sequences. The predicted local distance difference test (pLDDT) values above 70 correspond to a near-native structure prediction ^70,71^. The Predicted Template modelling (pTM) score and Interfacial pTM (ipTM) threshold were approximated to 0.3 and above^72^. These parameters and a high Predicted aligned error (PAE) score suggested possible interaction in our case. We report that Syt1 shows an interaction with the AP-3m1 with a pTM of 0.439, ipTM of 0.425, and a high mean pLDDT of 73.7 (Supplementary Fig 6B). Overall, a higher PAE score further indicates a possible interaction among Syt1 and AP3m1 subunit via a Tyrosine motif (YNSTG) at the C-terminal of Syt1. Meanwhile, we identified a Di-leucine motif EEKRAIL at the C-terminal of VMAT1, which is now predicted to interact with the AP-3s1 with pLDDT of 73.2, pTM of 0.63 and ipTM of 0.66 along with higher PAE score (Supplementary Fig 6C). Dlk1 also showed promising interaction with the AP-3s2 subunit at the C-terminal of Dlk1 based on a high PAE score (Supplementary Fig 6D). The tyrosine and di-leucine sorting motifs are important to direct cargoes to endosomal and lysosomal compartments from the TGN and Endosome or both^11,73,74^. These motifs have also been implicated in sorting Cargo by AP-1 complex via the σ and µ subunits, similar to what we see in the case of AP-3 as well^75^.

These interactions were validated in vivo in PC12 cells by a proximity ligation assay^76^ with Syt1, VMAT1 (where wild-type PC12 cells overexpressing VMAT-1 GFP probed using a GFP Ab), Dlk1, and AP3δ, which further showed a positive PLA signal, indicating a possible interaction between these cargoes and the AP-3 complex(Supplementary Fig 6E-H). Additionally, we have also performed co-immunoprecipitation experiments, further substantiating these physical associations with the AP-3 complex. In PC12 cells, Dlk1 was able to co-immunoprecipitate endogenous AP-3δ subunit (Supplementary Fig. 6i). Alternatively, the AP-3μ subunit was successful in co-immunoprecipitating the cargo VMAT1 (Supplementary Fig. 6J). Ultimately, AP-3 could be attributed to physical interactions with preferential DCV membrane cargoes Dlk1, VMAT1 and Syt1, which would be sorted to the DCVs. At the same time, depletion of AP-3 would allow these cargoes to be rerouted to the lysosomes for degradation, leading to an overall disruption of membrane trafficking functions and a defective regulated exocytosis.

## Discussion

The *C.elegans* AP-3 null worm gave the ideal starting point indicating a defective regulated exocytosis(Fig 1A-F), which directed us to investigate deeper into the roles of the AP-3 complex. Establishing an inducible AP-3 knockdown system in PC12 cells has been valuable in this study, elucidating new functional associations of AP-3 in the trafficking of regulated secretory vesicles and related organellar dynamics. Firstly, we show a clear impediment in the release of CgB upon stimulation (Fig 2B – G) that is further evidenced by a similar hindrance in the release of NPY (Fig 2h-i, Fig 3B-G). Although this impediment has been reported^19,21^, the conventional explanation accepted view hints at the role of the AP-3 complex in DCV biogenesis^19^. Therefore, this study focuses on the mechanistic insights and extensive mapping of the role of the AP-3 complex in regulating DCV function at multiple levels. We hypothesized AP-3’s involvement in the maturation of DCVs. The ultrastructural analysis denotes an increased DC size and decrease in the number of DCVs (Fig 5A and E, Supplementary Fig 2D and E), proximal positioning of DCVs next to the TGN (Fig 5F and G, Supplementary Fig 2G and H), this consistently indicates a maturation defect^45^. Accumulation of sPHM and CgB in the cells also points towards a defect in the maturation (Fig 2B-F and J, Supplementary 1A-B and K). Maturation also depends on the fusion of immature vesicles by Stx6-mediated homotypic fusion^46^. Although AP complexes may have redundant functions^47^, we report that loss of AP-3 prompts defective retrograde trafficking of stx-6 to the TGN (Fig 5H-i and Supplementary Fig. 2I-J) resulting in improper homotypic fusion of the immature vesicles. A similar sorting of stx-6 dependent on AP-1 and clathrin^46^, along with the VEF western blot for Stx-6 showing an increase in the clathrin-coated vesicle fraction also points towards AP-3’s role in trafficking stx-6 that we plan to investigate in the future separately. It is also noteworthy that the bigger size of the dense core might be a result of the accumulation of secretory cargo in dense cores of DCVs, leading to impaired regulated exocytosis caused potentially by fusion formation defects. Previous research has indicated the role AP-3 plays in post-Golgi trafficking^77^. We believe AP-3, despite its limited presence at the Golgi^78^, plays an important role in DCV and SLV exocytosis. Besides this, AP-3 also regulated Golgi integrity, and to the best of our knowledge, we are the first to report the effect of AP-3 on Golgi integrity. A similar Golgi reassembly defect does exist when a clathrin-heavy chain is lost^51^, which in turn suggests that the loss of AP-3 disrupts sorting sites on the TGN, which may cause a probable missorting of DCV cargoes at the TGN, leading to the rerouting of various cargoes.

At the endosomes, AP-3 has been implicated in forming synaptic vesicles and recycling exocytotic vesicles. Hence, the observed defect in the Endolysosomal processing of 10kDa dextran in the absence of AP-3 reveals the importance of AP-3 in the clearance via the Endolysosomal pathway. Also, AP-3 regulates Cargo sorting at the endosomes into the SLVs or lysosomes^63^. Since SLV bud from endosomes causes subsequent defects, we believe that DCV exocytosis is affected by AP-3. Our research indicates that defects in AP-3 cause budding deficiency at the Golgi (Fig 6) and, as such, cause mislocalization of several proteins that are significant to DCV exocytosis. We all show that AP-3 plays a definite role in DCV exocytosis through post-Golgi trafficking, showing an underlying mechanism, as seen earlier. In our investigation, we utilized proteomics to successfully identify novel cargo molecules, specifically Syt1, Dlk1, and VMAT1, that rely on AP-3 for proper sorting into dense core vesicles (DCVs). This finding shed new light on the intricate mechanisms governing DCV function.

Previous research by ^21^ reported the disappearance of Syt1 from DCVs in cells lacking AP-3; with our alphafold modelling study, we report a physical association of Syt1 with the m1 subunit of AP-3(Supplementary Fig 6B, 6F) via a tyrosine motif^11^. Our study serves as a valuable confirmation of this dependence, employing proteomic analysis and immunofluorescence techniques to strengthen the evidence. We also showed the presence of a Di-leucine motif^4^ at the C-terminal of VMAT-1, which is predicted to interact with the s1 subunit of AP-3 (Supplementary Fig 6C, 6G). The PLA and co-immunoprecipitation experiments in PC12 cells further validated this interaction between AP-3 complex (AP3δ subunit) and VMAT1 (Supplementary Fig 6J), indicating the dependence on VMAT1 on the AP-3 complex for sorting into the DCVs. However, identifying Dlk1 as an AP-3-dependent DCV cargo uniquely contributes to the understanding of regulated exocytosis of DCVs. Although we had seen weak physical interactions of the σ subunit of AP-3 with Dlk1 via the AlphaFold modelling due to the poorly resolved structure of Dlk1, invivo PLA and co-immunoprecipitation experiments in PC12 cells indicate physical interactions between Dlk1 and AP-3 complex (δ subunit) (Supplementary Fig 6D,H,I).

Dlk1 stands out as a completely novel discovery, highlighting the potential for AP-3 to interact with a wider range of molecules than previously thought. We are currently working on characterizing and validating these predicted physical interactors. There might be more novel candidates in our 66 proteins identified in the proteomics, which might be AP-3 dependent cargo, which becomes an interesting path to explore, and we plan on investing our time in them and gaining a comprehensive understanding of the overall functions associated with the AP-3 complex. We have proposed a model for the multifaceted role of AP-3 in the trafficking of DCV components and other membrane proteins from Golgi to their target. The significance of this research extends beyond these specific cargo identifications. By delving into AP-3’s function within DCVs, we contribute significantly to the overall understanding of DCV biology. Our findings help bridge knowledge gaps and establish novel functionalities for AP-3 within the cell, solidifying the hypothesis that AP-3 plays a multifaceted role in various cellular processes. In essence, this study paves the way for a more comprehensive picture of AP-3’s intricate role in regulating the subcellular transport of secretory vesicles and, thus, regulating specialized trafficking pathways(Supplementary Fig 7).

## Supporting information

no link

## Acknowledgements

The authors thank Yulong Li’s Lab, Tom Carter’s Lab, Ed Chapman’s Lab, Gary Miller’s Lab, John Paul Luzio’s Lab, and Cedric S. Asensio’s Lab for the various plasmids. The authors also thank Scottie Robinson and Subbarao for the kind gift of the sweet pea and mouse monoclonal antibodies, respectively, against the AP-3 delta subunit. The authors also thank the members of the B.S.S lab and Sandhya Koushika for critical feedback on the manuscript. Services of CIMR-Mass Spec facilities and support of Robin Antrobus are acknowledged. The authors also thank the DBT-Ramalingaswami fellowship, ICGEB research grant, IBRO for a research grant, SERB for a startup research grant and DBT-NBRC for providing valuable funds.

## Author Contributions

Conceptualization, B.S.S; Methodology, B.S.S, S.S and V.G; Investigation, S.S., V.G, C.M., B.S.C, S.B and B.S.S.; Writing – Original Draft, S.S., V.G and B.S.S.; Writing – Review & Editing, S.S. and B.S.S.; Funding Acquisition, B.S.S.; Resources, B.S.S; Supervision, B.S.S.

## Materials and Methods

### Cell Culture

PC12 cells (from ATCC) were maintained in DMEM media (Gibco, 12100046) supplemented with 10% Horse Serum (Gibco, 16050114), 5% Fetal Bovine Serum (Gibco, F7524) and Antibiotic/Antimycotic (Gibco, 15240062) in a humidified cell culture chamber at 37°C with 5% CO_2_. The cells were routinely treated with Mycoplasma removal agent (MP Biomedical, 30-50044) and used in experiments for no more than 9 passages. The SMARTvector rat lentiviral system (Dharmacon) was used to generate inducible shRNA-expressing cells that targeted the knockdown of Ap3δ and coexpressed GFP. The following constructs were used in the study: AP3sh1, 5’ – TGGTCAAGGGCAGTATCGA – 3’; AP3sh2, 5’ – GCTGAGAATCCGTACATTA – 3’; AP3sh3, 5’ – GAAGGTACTGACGTCATCA – 3’.The generation of PC12 cells stably expressing the inducible shRNA constructs, for future reference, AP3sh1, APsh2, and AP3sh3, were performed as described previously^24^. Briefly, lentiviral transduction containing the virions encoding three different constructs, as mentioned previously, was carried out in PC12 cells in poly L lysine (Sigma-Aldrich, P2636) coated 6 well plates according to the manufacturer’s protocol (Dharmacon). The cells containing the integrated lentiviral constructs were selected in 2 μg/ml puromycin (Sigma, 118M4079V), constantly replacing the medium for 3 days until resistant colonies were seen. The cells were induced with 5 μg/ml doxycycline (Himedia, 0000-348412) overnight, and highly expressing GFP-positive cells were obtained by fluorescence-associated cell sorting (FACS). The cells were maintained in a medium containing 0.2 μg/ml puromycin without doxycycline.

For experiments, AP3sh1, AP3sh2, and AP3sh3 cells were split and maintained in PC12 media containing 0.2 μg/ml puromycin and 5 μg/ml doxycycline for 5 days to induce the knockdown of AP3, labelled as +DOX or KD. Simultaneous maintenance of non-induced cells, labelled as -DOX or WT, was used as a control for each experiment.

### Mammalian expression constructs

All the expression plasmids used in this study

| NAME | MAKE | Catalog No. |
| --- | --- | --- |
| Synaptophysin-pHTomato | A kind gift from Yulong Li Lab, China | N/A |
| NPY-pHTomato | Addgene | 83501 |
| NPY-mApple | Addgene | 83498 |
| mCherry-Lysosomes-20 | Addgene | 55073 |
| VMAT1-mCherryN1, VMAT1-EGFPN1 | A kind gift from Tom Carter's Lab, London | N/A |
| VMAT2-mCherryN1 | A kind gift from Gary Miller's Lab, US | N/A |
| Syt1-mCherryN1 | A kind gift from Ed Chapman's lab, US | N/A |
| Syt4-mCherryN1 | A kind gift from Ed Chapman's lab, US | N/A |
| Syt7-mCherryN1 | A kind gift from Ed Chapman's lab, US | N/A |
| Phogrin-mCherryN1 | A kind gift from John Paul Luzio's Lab, UK | N/A |
| NPY-mCherry-SBP | A kind gift from Cedric S. Asensio's Lab, US | N/A |
| stim/sp-str-linker-kdel-hook | A kind gift from Cedric S. Asensio's Lab, US | N/A |

### Transfections, Immunofluorescence and Confocal Imaging

AP3sh1 and AP3sh2 were transfected using METAFECTENE® PRO (Biointex, Lot. No. RKP205/RK081621) through the protocol provided by the manufacturer^79^. In summary, plasmid DNA and METAFECTENE® PRO were mixed separately in DMEM medium without any supplements, mixed and incubated for 20 minutes at room temperature for complex formation. The complex was added directly to the cells grown in DMEM with 10% horse serum and 5% fetal bovine serum within 4 hours of cell seeding. For stable transfection of cells, they were seeded in a 12-well plate and transfected according to the protocol described. Upon 12 hours the cells were incubated in PC12 media for 24 hours. The transfected cells were then transferred to a 60mm dish and cultured in PC12 media containing 2 mg/ml G418 (GoldBio, G-418-1) for 14 days in order to select cells containing NPY mApple and NPY pHTomato stably.

For transient transfections to see the intracellular localization, WT and KD cells were seeded on a 15mm coverslip (Assistent/ Glaswarenfabrik). As mentioned previously, a complex containing the plasmid and METAFECTENE® PRO were added directly to each coverslip within 4 hours of seeding. The cells were kept in PC12 media 12 hours post seeding for at most 72 hours after which the cells were utilized for Immunofluorescence or live cell imaging. The transfections carried out were of the plasmids, Syt1 mCherry, Syt4 mCherry, Syt7 mCherry, VMAT1 mCherry, VMAT2 mCherry, Phogrin mCherry, Lyso20 mCherry.

For Immunofluorescence, a previously standardized protocol was followed^80^. In brief, WT and KD cells were seeded onto PLL coated 12mm coverslips. Cells were rinsed with 1X PBS (Gibco, 21600-069) and fixed with 4% paraformaldehyde (PFA)(Fisher Scientific, 30525-89-4) for 20 minutes at room temperature. Post fixation, cells were again gently rinsed with 1X PBS (3 times for 5 minutes each) and then permeabilized using 0.1% Triton X-100 (Sigma Aldrich, 9002-93-1) for 10 minutes. The permeabilized cells were again rinsed with 1X PBS (2 times for 5 minutes each) and blocked using 1% bovine serum albumin (BSA)(Sigma, A4503) for 1 hour. Following that, the cells were subjected to incubation with a primary antibody overnight. Post incubation with primary antibody, cells were rinsed again with 1X PBS (3 times for 5 minutes each) and incubated in secondary antibodies(Invitrogen Rabbit Alexa Fluor™ 647(A-31573), Mouse Alexa Fluor™ 647(A-31571), Rabbit Alexa Fluor™ 594(A-11012), Mouse Alexa Fluor™ 594(A-11005)), with their cognate primary antibodies, followed by gently washed with 1X PBS (3 times for 5 minutes each). The coverslips were then mounted using Flouroshield^TM^ (Sigma, F6057) to reduce photobleaching. All images were captured using a Nikon A1HD25 confocal microscope under a 60X oil objective (NA 1.4) at a 2048*2048-pixel resolution unless mentioned differently.

### Immunoblotting

WT and KD cells were lysed using RIPA buffer (10 mM Tris-HCl (Sisco Research Laboratory Pvt. Limited, 37969/ 77-86-1), 150 mM NaCl (MP Biomedicals, 194848), 1 mM NaVO4 (Sigma Aldrich, S-6508), 30 mM Na4P2O7 (Sigma Aldrich, 011K0307), 50 mM NaF (Sigma Aldrich, 7681-49-4), 1% Nonidet P-40 (Sigma Aldrich), 0.1% SDS (Affymetrix USB Products, 151-21-3), 1 mM PMSF (Sigma Aldrich, 52K0052), 1% Triton X-100 (Sigma Aldrich), 0.5% C24H39NaO4 (Sigma Aldrich, 145224-92-6), and dissolved protease inhibitor cocktail (Roche Diagnostics, 11714900) in water, pH 7.4). The protein contents of the cell lysate were quantified using a BCA Protein Assay Kit (TAKARA) following the manufacturer’s protocol. ^81^ The samples were separated using a 10% SDS-polyacrylamide gel and transferred onto nitrocellulose membranes. After blocking with 5% BSA (Sigma Aldrich), membranes were incubated with primary antibodies, followed by incubation with secondary antibodies (Peroxidase AffiniPure Goat Anti-Rabbit IgG (H+L) Jackson Immuno Research 111-035-003, Peroxidase AffiniPure Goat Anti-Mouse IgG (H+L) Jackson Immuno Research 115-035-003). The membrane was washed thrice for 10 minutes after primary and secondary antibody incubation to remove excess antibodies. ECL was prepared using the previously described protocol, and bands were visualized using the Uvitec Mini HD9 gel imaging system ^82^. Densitometry quantifications were done using ImageLab (6.0.1) ^83^.

### Stimulus Coupled Secretion Assay

A stimulus-coupled secretion assay was performed in PC12 cells according to a previous protocol ^24,40^. In brief, WT and KD cells were plated in a 60mm dish (Tarson) at a confluency of 90%. 24 hours post cell seeding, WT, and KD cells were incubated with basal buffer (150 mM NaCl, 5 mM KCl, 2 mM CaCl_2_, 10 mM HEPES; pH 7.4) and stimulation buffer (150 mM NaCl, 100 mM KCl, 5 mM BaCl_2_, 10 mM HEPES; pH 7.4) for 20 minutes at 37°C in a humidified cell culture chamber with 5% CO_2_. The supernatant was then collected, whereas cell lysates were prepared, as mentioned previously, for Immunoblotting. Supernatants and cell lysates were collected and run in SDS - PAGE and probed with CgB (Proteintech, 14968-1-AP,1:5000), Dlk1(Proteintech,10636-1-AP), GAPDH(Proteintech,60004-1-Ig,1:10000) and B-Tub (Proteintech,10094-1-AP,1:20000) for AP3sh1, AP3sh2 and AP3sh3.

A stimulus-coupled secretion assay was also performed using a different secretagogue to assess regulated secretion using KCl^40^. In brief, WT and KD cells were seeded, as previously mentioned, to perform a secretion assay. The cells were incubated with basal buffer (150 mM NaCl, 5 mM KCl, 2 mM CaCl2, 10 mM HEPES; pH 7.4) and stimulation buffer (55 mM NaCl, 100 mM KCl, 2 mM CaCl2, 10 mM HEPES; pH 7.4) for 20 minutes and process as mentioned previously. Supernatants and processed cell lysates were subjected to SDS-PAGE and probed further with CgB and JH610(sPHM) (1:2000) in AP3sh1 and AP3sh2 cell lines.

Stably selected cells of NPY-mApple in an AP3sh1 and AP3sh2 background were also subjected to a stimulus-coupled secretion assay^80^. In summary, WT and KD cells harboring NPY-mApple were seeded in a 6-well plate at 90% confluency and incubated with basal buffer (150 mM NaCl, 5 mM KCl, 2 mM CaCl2, 10 mM HEPES; pH 7.4) and stimulation buffer (55 mM NaCl, 100 mM KCl, 2 mM CaCl2, 10 mM HEPES; pH 7.4) for 20 minutes at 37°C in a humidified cell culture chamber with 5% CO_2_. Collected supernatants and cell lysates were processed as mentioned previously. For the plate reader assay, 10μl of the cell lysate and the supernatants were diluted at 1:10 in 1X PBS (Gibco). 100μl of the solution was loaded in a dark 96 well plate (Tarson) in duplicates along with subsequent controls for background subtraction. NPY-mApple fluorescent signals were then recorded using the TECAN plate reader (Ex λ-568 nm, Em λ-592 nm), subtracting them from the autofluorescence of the plate itself ^84^.

### Constitutive Secretion

To check the constitutive release of proteins in basal conditions, secreted fractions from unstimulated cells were analyzed by SimplyBlueTM SafeStain (Invitrogen, LC6060) Coomassie staining ^39,40^. In brief, WT and KD cells were seeded in 60 mm culture dishes at a density of 10^6^ per dish. The cells were incubated with basal buffer (150 mM NaCl, 5 mM KCl, 2 mM CaCl2, 10 mM HEPES; pH 7.4) for 3 hours, during which the supernatants and cell lysates were collected and processed as mentioned previously in the report. The samples were resolved using a 10% SDS-polyacrylamide gel. The gel was washed with Milli-Q water twice and then stained with SimplyBlueTM SafeStain (Invitrogen) for 1 hour at room temperature with gentle shaking. To remove the background, the gel was washed with Milli-Q water twice and later imaged in the Azure c300 gel imaging system, and all the automatically detected bands were quantified using ImageLab (6.0.1). Furthermore, for estimating constitutive secretion using a marker, Immunoblotting was carried out and probed for HSP90 (Proteintech,13171-1-AP,1:1000). Bands were visualized using the Uvitec Mini HD9 gel imaging system ^82^.

### Live Cell Imaging and Analysis

Stably selected cells of NPY-pHTomato in an AP3sh1 and AP3sh2 background were subjected to live cell imaging. Synaptophysin pHTomato (Addgene) was transiently transfected in AP3sh1 cells^85^. For imaging, WT and KD cells were plated on 15mm coverslips. On the day of the experiment, cells were rinsed with basal buffer (150 mM NaCl, 5 mM KCl, 2 mM CaCl2, 10 mM HEPES; pH 7.4), and coverslips were placed in an open imaging chamber (Life Cell Instruments, Korea). The images were captured using Nikon Eclipse Ti2 at 100X oil (1.30) at 300 ms for 2 sec in the Tyrode& buffer (basal), after which stimulation was given at 300 ms for 15 sec in the Tyrode’s buffer (basal). At the end of the experiments, Tyrode’s solution containing 100 mM NH4Cl, pH 7.4, was added to its final concentration, and images were taken to acquire the total fluorescence of the transfected cell. Captured time-lapse movies were analyzed using ImageJ software, and after background subtraction, the change in puncta intensity/cell (Δf) after 100 mM KCl stimulation compared to basal fluorescence (f_0_) was plotted as Δf/f_0_.

### Transmission Electron Microscopy

Sample preparation for transmission electron microscopy was carried out according to a previous protocol.^80^ Briefly, WT and KD cells from AP3sh1 and AP3sh2 were seeded in a 100mm dish. The cells were fixed overnight with routine fixative, 2% paraformaldehyde (PFA) (Fisher Scientific) and 2.5% glutaraldehyde (Sigma – Aldrich, MKBG0637V) in 0.15M sodium cacodylate buffer (Sigma – Aldrich, C0250). Cells were then pelleted at 10,000 rpm for 10 minutes in fresh fixative. Post-fixation was done for 1 hour in 1% osmium tetroxide (Sigma, 05500), followed by three washes with 0.15M sodium cacodylate buffer. Subsequent dehydration was done through graded ethanol incubations (10 minutes each of 50%, 70%, 90%, 2 x 100%). Pelleted cells were then incubated in propylene oxide (Tokyo Chemical Industry, E0016) (2 x 20 minutes), followed by overnight infiltration in a 1:1 mixture of propylene oxide and Araldite 502 (Ted Pella, 18060). The samples were subsequently embedded in a mixture of Araldite 502, DDSA (Ted Pella, 18022), and DMP-30 (Ted Pella, 18042) as per a previous protocol ^86^ and polymerized at 60°C for 72 hours. Ultrathin sections were cut on a Leica UCT ultramicrotome, picked onto carbon-coated copper grids (Electron Microscopy Sciences), stained with 10% Uranyl acetate (Sisco Research Laboratory, 81405) and counter-stained with Sato’s lead citrate (Electron Microscopy Sciences, 17800) and imaged using JEOL JEM1400-plus transmission electron microscope. The images were used for the morphometric analysis of dense core (DC) and dense core vesicles (DCV). Furthermore, intracellular DCV distribution was obtained by measuring the shortest distance of the DCV from the cell membrane.

### Retention Using Selective Hook Assay and Analysis

1.5 million Cells were seeded in each of two 60 mm dishes with 7 coverslips. After 4 hours, cells were transfected with NPY-SBP-MCherry (Cargo) and KDEL-Str (Hook) plasmids (a kind gift from CS Asensio lab) in a 1:3 ratio ^49^. After 8 hours, transfection media was replaced with full media, and cells were grown for 48 hours before the experiment. 50 mM Biotin(Sigma, B4501-1G) to the final concentration in Opti-Mem was used to replace the full DMEM for the experiment, and coverslips from each condition were fixed at 5 different time points (0 Mins, 20 Mins, 40 Mins, 60 Mins, and 90 Mins) post Biotin addition ^87^. Then, fixed coverslips were subjected to Immunofluorescence staining for Golgin 97(Proteintech,12640-1-AP, 1:1000) and visualized using a Confocal Microscope as described previously. For the analysis, Mander’s coefficient tM1 was used to measure the colocalization between Golgin 97(Golga1) and Cargo (tagged with mCherry) for all the time points. To measure the Rate of entry and the Rate of exit of Cargo at Golgi, the slope was calculated as, Rate of entry= <u>(tM1 at 20 mins) -(tM1 at 0 mins)</u> / 20-0 Rate of Exit= <u>(tM1 at 40 mins) -(tM1 at 20 mins</u>) / 40-20 Where tM1 corresponds to the colocalization mander’s coefficient between Golgi and NPY mCherry. This entry and exit rate were further normalized to the average rate of entry and exit in MOCK cells.

### Brefeldin A Assay

In a 12-well plate with 15mm coverslips, 200,000 cells were seeded using the WT(-DOX) and KD cells. After 24 hours, the cells were treated with 2.5ug/ml of BFA in DMEM complete media (10%HS+5%FBS) for 15 mins at 37 degrees Celsius inside a CO2 incubator. The first coverslip was washed with 1X PBS and fixed with 4% PFA before the treatment with BFA (Sigma, B6542) to serve as the control for the experiment. After the BFA treatment, cells were washed with 1X PBS twice, and the second coverslip was fixed with 4% PFA, while the other coverslips were incubated with DMEM complete media without any BFA (this is called the washout step). Further, the coverslips were fixed at 20 minutes and 40 mins post-washout, respectively. Once all the coverslips were fixed, Immunofluorescence was done using a Golgin 97 antibody to visualize the Trans Golgi network using a confocal microscope. This protocol was adapted from^88^. For analysis, the puncta visible in the images were counted using the analyze particle plugin of image J and were normalized per cell^89^. Multiple t-tests were performed for each condtiton.

### SILAC, vesicle-enriched fraction preparation, Immunoblotting, and mass spectrometric analysis

For Stable Isotope Labelling by Amino Acids in Cell Culture (SILAC) experiments, PC12 cells were cultured for at least 3 weeks to achieve metabolic labelling in “SILAC medium”. The “SILAC medium” consisted of DMEM (Silantes) supplemented with 5% (vol/vol) dialyzed Fetal Bovine Serum (Silantes) and 10% (vol/vol) Horse Serum (Silantes) and either “heavy” amino acid (l-arginine - ^13^C_6_ ^15^N_4_ (50 mg/l) and l-lysine - ^13^C_6_ ^15^N_2_ (100 mg/l)) (Silantes) or “light” amino acid (l-arginine - ^12^C_6_ ^14^N_4_ and l-lysine - ^12^C_6_ ^14^N_2_) (Sigma Aldrich). The average incorporation efficiency was calculated to be ∼98%, as seen by mass spectrometry.

For vesicle-enriched fraction preparation, the protocol was followed as described previously ^24,55^. To summarize, 2 fully confluent 500 cm2 dishes (Corning) were seeded with WT or KD cells for each preparation. At 4°C, each dish was rinsed with 1X PBS and scraped with 12 ml of Buffer A (0.1 M MES, pH 6.5 [adjusted with NaOH], 0.2 mM EGTA and 0.5 mM MgCl2). The cells were then homogenized using a motorized Dounce homogenizer (25 strokes) and passed through a syringe using a 21G X 2” needle. 200 μL of the homogenized solution was kept separately and processed for cell lysate, whereas the rest of the solution was centrifuged at 4500 rpm for 30 minutes. The supernatant was treated with 50 μg/ml ribonuclease A (MP Biomedical) for 60 min at 4°C. The digested ribosomes were pelleted by centrifugation at 4500 rpm for 15 minutes. The resulting supernatant was centrifuged at 55,000 rpm for 45 minutes (MLA 80 rotor, Beckman Coulter). The pellet was resuspended in 350 μl of Buffer A and again subjected to homogenization using a 1 ml Dounce homogenizer (69 strokes) and mixed with an equal volume of FS buffer (12.5% [wt/vol] Ficoll, 12.5% [wt/vol] sucrose in Buffer A). The solution was then pelleted, and the supernatant was diluted 5 times in Buffer A and centrifuged at 35,000 rpm for 35 minutes (TLA-110 rotor, Beckman Coulter). The pellet was then identified as the vesicle-enriched fraction (VEF) and processed accordingly for mass spectrometry and Immunoblotting.

For Immunoblotting, AP3sh1 and AP3sh2 cells were used. In brief, both WT and KD cells were seeded in 2X 500 cm^2^ dishes and subjected to vesicle-enriched fraction preparation, as mentioned above. 2X non-reducing SDS buffer (4% [wt/vol] SDS and 10 mM Tris-HCl, pH 8.0) ^90^ was mixed in a 1:1 ratio with the cell lysate, and the subsequent solution was passed through the QIA shredder (Qiagen). For the VEF, 50 μl 1X non-reducing SDS buffer was used for resuspending the pellet. The samples were heated at 95^0^C for 5 minutes and processed, as mentioned before, for Immunoblotting. In all cases, 20 μg of cell lysate was loaded, whereas 2.5 μg of VEF was loaded. The blots were processed accordingly and probed for targeted proteins CgB, Clathrin Heavy chain (CHC, Proteintech 26523-1-AP,1:2000), Syt1(Synaptic systems 105011,1:2500), ZnT3(Proteintech 17363-1-AP, 1:2500), Syntaxin 6(Synaptic systems 110062, 1:2000), VMAT1(Proteintech 20340-1-AP, 1:2000), AP3m1(Proteintech 12114-1-AP, 1:2000), Dlk1(Santa Cruz SC-25437,1:500). For mass spectrometry, metabolically “heavy” and “light” cells were cultured accordingly. For the first experiment in AP3sh1, WT cells were cultured in a “heavy” medium, whereas KD cells were cultured in a “light” medium. In the second experiment, WT cells were cultured in a “light” medium, whereas KD cells were cultured in a “heavy” medium. The VEFs from each condition were resuspended in 30 μl of 1X non-reducing SDS buffer and then mixed. The samples were run in a NUPAGE precast 12% gel (Invitrogen) and stained with SimpleSafe Blue Stain (Thermo Fisher). Each lane in the gel was cut into eight pieces and then analyzed for mass spectrometry. The resolved proteins were reduced, alkylated, and digested in-gel with trypsin, and then the resulting peptides were concentrated and desalted using Stage tips^91^. Tryptic peptides were analyzed by LC-MS/MS using a Q-Exactive coupled to an RSLCnano3000 UHPLC (Thermo Fisher). Peptides were resolved using a 50-cm EASY-Spray PepMap C18 column (Thermo Fisher) with data acquired in a data-dependent acquisition manner.

Data was processed using MaxQuant v2.1.4.0 using Andromeda to search a Uniprot *Rattus norvegicus* database (downloaded 7 April 2014, 27,344 entries) with *Homo sapiens* AP1G1 (O43747) added. N-terminal protein acetylation and methionine oxidation were set as variable modifications, carbamidomethyl cysteine was set as a fixed modification, and both LFQ and iBAQ were enabled. Oxidised methionine was set as variable modifications and carbamidomethyl cysteine as fixed. Default false discovery rate calculations were used as provided by MaxQuant. Where indicated, ratios were normalized on protein average. The raw data files were processed using MaxQuant (with requantify and match between runs features enabled). The primary output for each SILAC comparison was a list of identified proteins, a ratio of relative abundance (heavy/light ratio), and the number of quantification events (count). Each MaxQuant output file was formatted identically: reverse hits, proteins identified only by site, common contaminants, and proteins with no gene name were removed (refer to Supplementary Table 1).

### Dextran Uptake Assay

A previously published protocol was used to understand the modes of exocytosis ^73^. In brief, WT and KD cells were washed once with 1X PBS and twice with extracellular bath solution (145 mM NaCl, 2.8 mM KCl, 2 mM CaCl_2_ and 100 mM HEPES, pH 7.4) and then incubated for 5 minutes with 50uM dextran (10 and 40 kDa, Invitrogen D1817 and D1842) in 100 mM KCl containing extracellular bath solution at 37°C. The unbound dye was washed out with the previously mentioned bath solution (without Ca2+) and immediately fixed with 4% PFA. Confocal imaging was done, as previously mentioned, where a Z-series of 1μm optical sections were scanned. For analysis, the consecutive optical sections were projected in ImageJ, and the total numbers of dextran fluorescent puncta per cell were determined and normalized according to the area of the cell.

### Catecholamine uptake Assay

WT and KD cells, cultured in six-well plates coated with poly-l-lysine and reaching 80% confluence, were exposed to 1 µCi [3H] l-norepinephrine for a 3-hour incubation period. Subsequently, they underwent two washes with release buffer and were subjected to a 15- minute incubation at 37°C, either in release buffer or stimulation buffer. Following this, supernatants were collected, and the cells were lysed in a lysis buffer. Both the supernatants and lysates were combined with a liquid scintillation cocktail, and the radioactivity was measured using a Beckman liquid scintillation counter. The experiments in Figure 7 and Supplementary Fig 5 represent biological replicates conducted independently thrice. This protocol was adapted from ^24^.

### Zinquin Ethyl Ester Assay

To visualize the zinc uptake, we used a previously published protocol ^62^ using Zinquin ethyl ester(Abcam, GR241459-21) dye. WT and KD cells were seeded on 15mm coverslips in 12 well plates and kept overnight. The next day, cells were incubated with a solution of ZnSO4(SRL, 7446-20-0) containing 50uM ZnSO4, 10mM Sodium pyruvate (Gibco, 11360-070), and 10mM glucose (MP Biomedical, 194672) in HBSS (with no calcium and magnesium) for 1 hour at 37 degrees Celsius. Later, the cells were washed with 1X PBS and incubated with 50uM of Zinquin ethyl ester (in 10mM glucose in PBS) for 20 mins at 37 degrees Celsius.

After which, the cells were washed multiple times (minimum twice) with 1X PBS and were fixed with 4% PFA for 10mins. These coverslips were imaged in the Nikon Eclipse Ti2 at 100X oil (NA 1.30).

### Endolysosomal Assay

In this assay, 200,000 cells seeded on coverslips in a 12-well plate from both WT and KD cells were subjected to incubation at 37 degrees Celsius with 1% 10kDa Dextran (50um) in DMEM incomplete media (i.e., no serum and antibiotic) for 30 mins. After the cells were washed with 1X PBS twice, the individual coverslips were fixed at 0-, 30-, and 60-mins post-washing, respectively, with 4% PFA. Confocal imaging was done, as previously mentioned, where a Z-series of 1μm optical sections were scanned. For analysis, the consecutive optical sections were projected in ImageJ, and the total numbers of dextran fluorescent puncta per cell were determined and normalized according to the area of the cell.

### Protein-protein interaction using ColabFold

Protein sequences for AP-3 complex subunits, Syt1, Dlk1, and VMAT1 specific for rats were taken from UniProt. To run the modelling, the protein sequences were fed onto ColabFold^69^ without changing any default settings. After the modelling was completed with ColabFold, using the different algorithms, generated predictions ranked 1 to 5. Based on the different parameters, such as pLDDT, pTM, ipTM and PAE score, the most suitable interaction was selected. The PDB files generated from these algorithms were analyzed using chimeraX and the PyMOL software to identify the interacting amino acids. The interactions reported were thresholded to a bond length of 4 angstroms or lower, which included both non and non-polar contacts^92^.

### C. elegans maintenance, imaging, and analysis

All *Caenorhabditis elegans* strains were grown at 20°C on standard Nematode Growth Medium (NGM) agar plates seeded with *Escherichia coli* OP50 bacteria.^36^ The strains used in this study are KP3947 *nuIs183* [*unc-129p*::nlp-21::Venus + *myo-2p*::GFP], a kind gift from Dr. Prof. Kavita previously used in^35^ and RB662 *apb-3(ok429)* procured from the Caenorhabditis Genetics Centre (CGC; University of Minnesota, USA). *apb-3(ok429); nuIs183* was built using standard procedures. L4 stage animals were taken, anesthetized with 10 mM Sodium Azide, and mounted on a 5% agar pad. Images were acquired using a 60X/1.35 NA oil objective with 1 × 1 binning using Olympus IX83 fitted with a Yokogawa CSU-W1 excited with 488nm with 5% laser power and 300 ms exposure for wild type and mutant. Fiji-ImageJ ^93^ was used to generate representative images and analysis. For the dorsal nerve cord (DNC) fluorescence measurements, a maximum intensity projection was obtained from image stacks near the posterior dorsal nerve cord. The average intensity was calculated by drawing a segmented line along the DNC, and background intensity was subtracted by moving the same line adjacent to the DNC in the body of the worm. Maximum intensity projection was obtained from the image stack in the posterior region to analyze the intensity of the coelomocyte. The mean intensity and area covered by NLP-21 punctate above a certain threshold in the coelomocyte region were calculated. The threshold was maintained the same for wild type and mutant. All fluorescence values were normalized to wild-type controls for comparison. OriginPro 2020b was used to generate representative graphs and Statistical analyses.^94^ To test the normality of various distributions, the Shapiro–Wilk test was used. Statistical comparisons for normally distributed data employed the unpaired two-tailed Student’s t-test, while those for non-normally distributed data were conducted using the Mann–Whitney U test.

### Colocalization and Statistical Analysis

All images captured by Nikon A1HD25 confocal microscope under a 60X oil objective (NA 1.4) at a 2048*2048-pixel resolution were deconvoluted at the time of capture. Image J plugin JaCoP (Just another Colocalization plugin) was used for colocalization. The tM1 values from the colocalization were then converted to a percentage by multiplying the decimal values by 100 and represented at percentage colocalization. All quantitative analyses were performed in Prism 8.0. Graphs represented as mean ± SD were statistically compared using Student’s unpaired *t*-test (GraphPad Prism 8.0) when the data was shown to be normally distributed. For cases where the data did not pass the normality test, statistical comparisons between the two groups were done using the Mann-Whitney test (GraphPad Prism 8.0). Differences between groups were considered statistically significant for *p* values < 0.05. All the experimental data shown were analyzed from at least three independent experiments or at least eight cells.

### Proximity Ligation Assay

The PLA^76^ was carried out using the Duolink In Situ Red Starter Kit(Mouse/Rabbit DUO92101; Sigma-Aldrich) to assess the in vivo interaction among AP3δ and Syt1, AP3δ and VMAT1-GFP also, AP3δ and Dlk1 in WT PC12 cells. The cells were fixed in 4% PFA for 20 minutes and washed twice with PBS. 0.2% TritonX-100 in PBS was used to permeabilise cells. The primary antibody used was AP3δ (AB_2056641, Dilution 1:500), Dlk1(10636-1-AP, Dilution 1:500), Synaptotagmin 1(105011, Dilution 1:1000) and GFP (50430-2-AP, Dilution 1:500). The further protocol as followed as described^76^. All images were captured using a Nikon A1HD25 confocal microscope under a 60X oil objective (NA 1.4) at a 1024*1024-pixel resolution as Z stack images with a step size of 0.3 um. For representation, the images were Z-projected at maximum intensity. The PLA signal specifically corresponds to the red dots seen in the images. For negative control, we used a no-antibody control, which showed no PLA signal.

### Native Co-immunoprecipitation

We have used the soft elution method for native co-immunoprecipitation experiments as described^91^. The beads were washed four times with PBS-T (phosphate-buffered saline containing 1% Triton X-100), followed by a single wash with PBS to remove detergent. Bound proteins were eluted by incubating the beads in 100 µL of soft elution buffer (0.2% SDS, 0.1% Tween-20, 50 mM Tris-HCl, pH 8.0) at 25°C for 7 minutes with shaking. The elution was repeated once, after which the eluates were combined and centrifuged at 16,000 × g for 1 minute to remove residual beads. IP eluates and IgG IP samples were loaded on 10% SDS-PAGE gel and analysed by immunoblot. 1% of the initial homogenates was loaded as input.

## Supplemental information

**Document S1. Figures S1–S7**

**Table S1. Mass-Spectrometry Data**

**Fig. S1.**
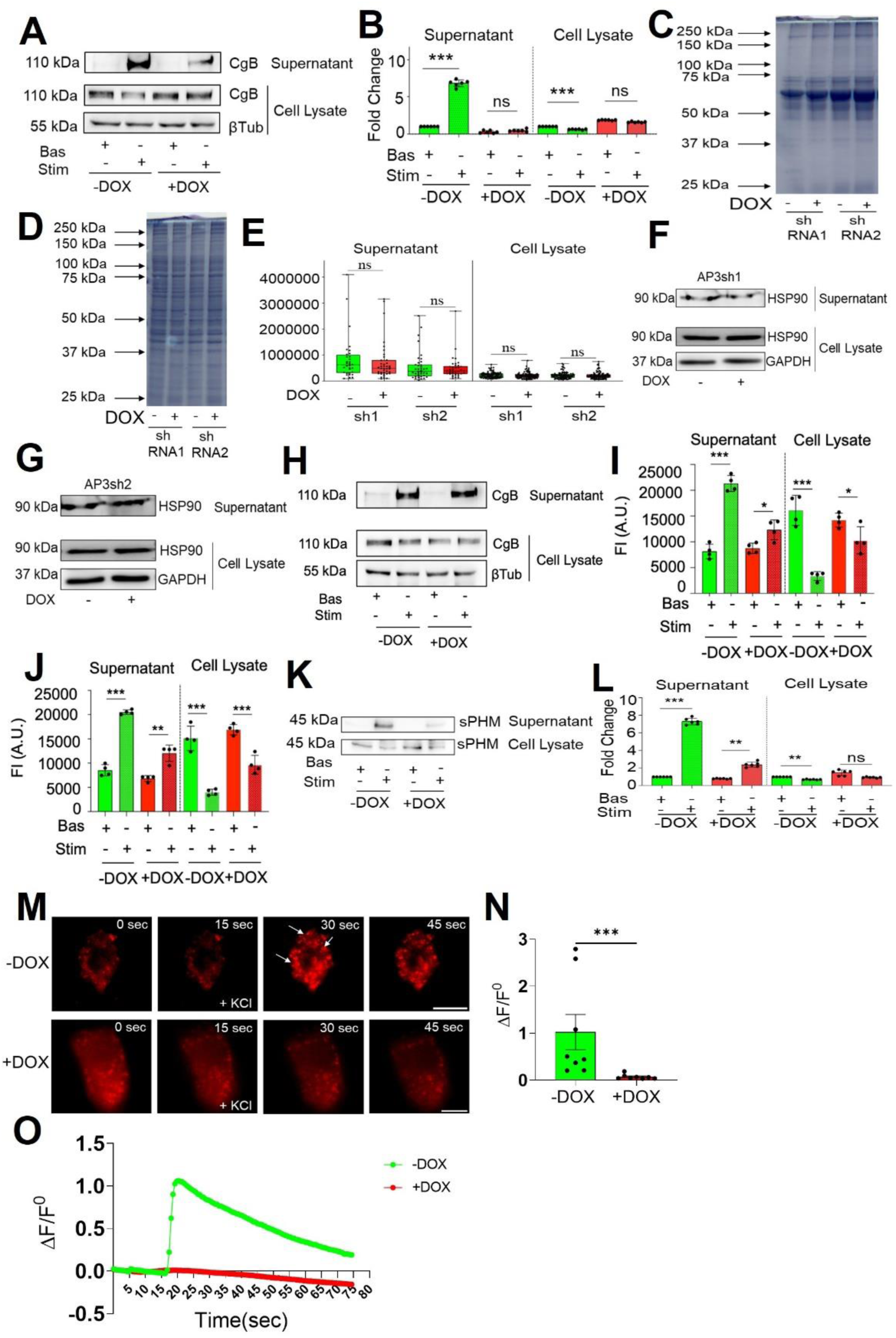
Stimulus-coupled secretion experiments and Live-Cell imaging reveal compromised DCV exocytosis in AP-3-depleted cells. Western blot and quantification (a-b) showing the secretion of CgB in mock-treated and AP-3 depleted cells (shRNA2) using BaCl_2_ as a secretagogue also indicate a similar defect in the induced release of CgB. n=3, Data shown as mean ± SD. (c-g) The gel picture shows the global constitutive signature and the corresponding quantification considering the auto-detected band intensities in Image-lab in shRNA1 and shRNA2 AP-3 depleted cells where each dot represents individual band intensities, n=3. Data is shown as mean ± SD. (f, g) Western blot showing no change in the constitutive release of HSP90 (protein unique to constitutive secretory pathway) in mock-treated and shRNA1 and shRNA2 AP-3 depleted cells. (h) The western blot shows no change in the case of CgB secretion for scrambled shRNA cells. (i,j) Fluorescent intensity was measured using a plate reader of mock-treated abd AP-3 depleted cells (shRNA1 and shRNA2) transfected with NPY-mApple,n=3 showed depletion in NPY release. (k-l) The release of sPHM from mock-treated (Green) and shRNA1 AP-3 depleted cells (Red) upon stimulation with a secretagogue KCl, (k-l) WB, and quantification show a major defect in release (n=3) (*P<0.05, **P<0.01, ***P<0.001; n.s. statistically non-significant). (m) Representative images of mock-treated and shRNA2 AP-3 depleted cells transfected with NPY-pHtomato (red) showing basal and stimulated conditions. Scale bar 5um (n) Quantitative analysis of the maximum Δf/f0 in mock-treated (n=8 cells) and shRNA2 AP-3 depleted cells (n=8 cells). (Mann-Whitney Test). (o) Representative time traces of mean NPY-pHtomato fluorescence intensities showing an average of all puncta per cell (green, mock-treated; red, shRNA2 AP-3 depleted cells). (N=3). (‘N’ denotes the number of independent experiments, and ‘n’ is the number of cells taken for quantification) Data are shown as mean ± SD. (*P<0.05, **P<0.01, ***P<0.001; n.s. statistically non-significant).

**Fig. S2.**
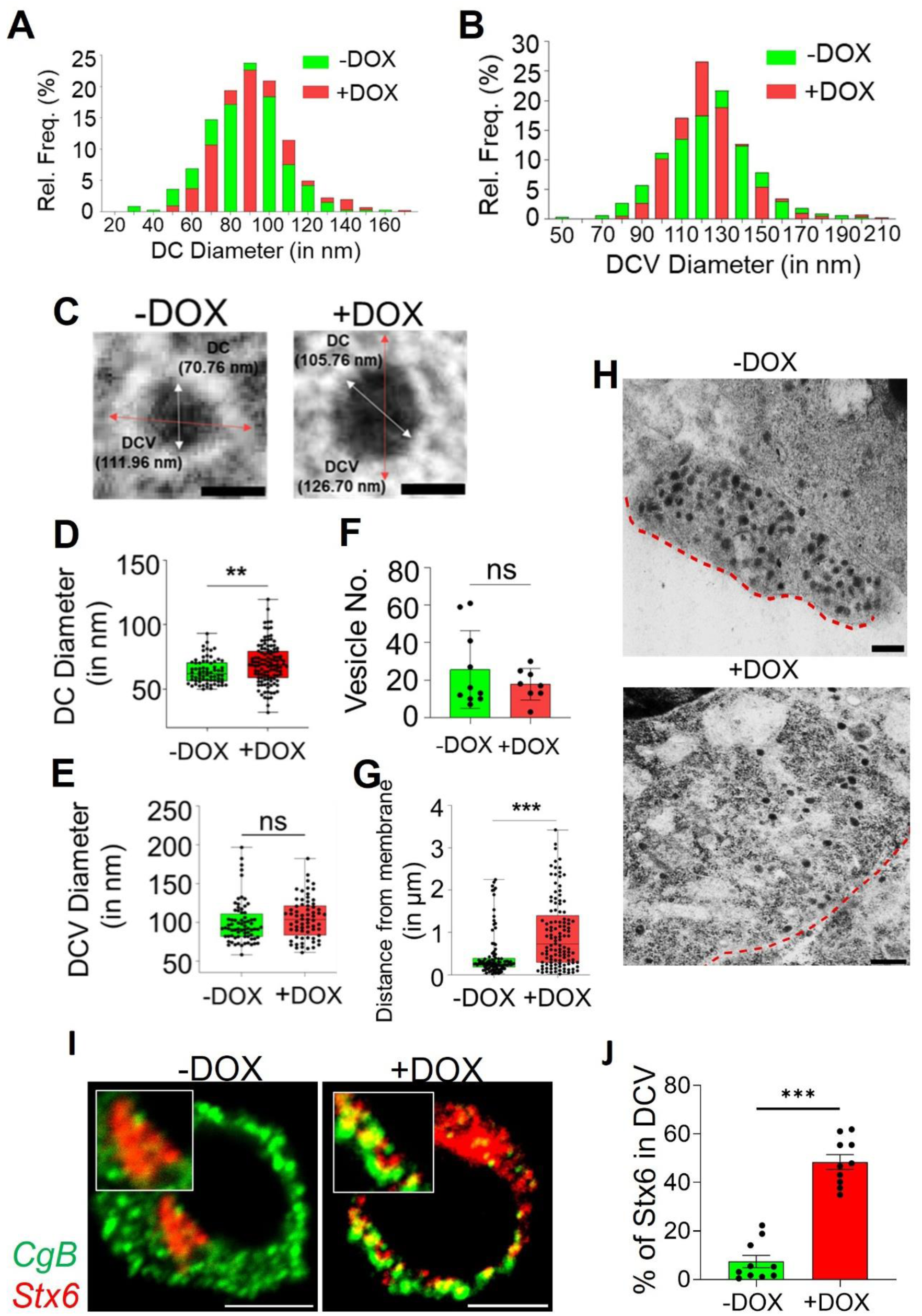
AP-3 affects the maturation of DCVs in PC12 cells. (a-b) Graphs for the relative frequency distribution of the DCVs with respect to DC diameter and DCV Diameter from the electron microscopy images for mock treated (green) and shRNA1 AP-3 depleted cells (Red). (c) Representative images showing DCVs for Mock treated and shRNA2 AP-3 depleted cells. Scale bar: 100nm. Graphs showing an increase in DC diameter (d), unaffected DCV diameter (e), as well as a graph for the number of DCVs (f). (g-h) Representative images and graph showing DCVs and their distance from plasma membrane increasing farther in case of AP-3 KD. (i-j) Representative confocal images and quantification showing an increase in colocalization (% colocalization) of Stx6 (Red) with CgB (DCV marker) (N=3, n=14 cells each) (Green). Scale bar: 200nm. (n=9-10 cells for each condition *P<0.05, **P<0.01, ***P<0.001; n.s. statistically non-significant).

**Fig. S3.**
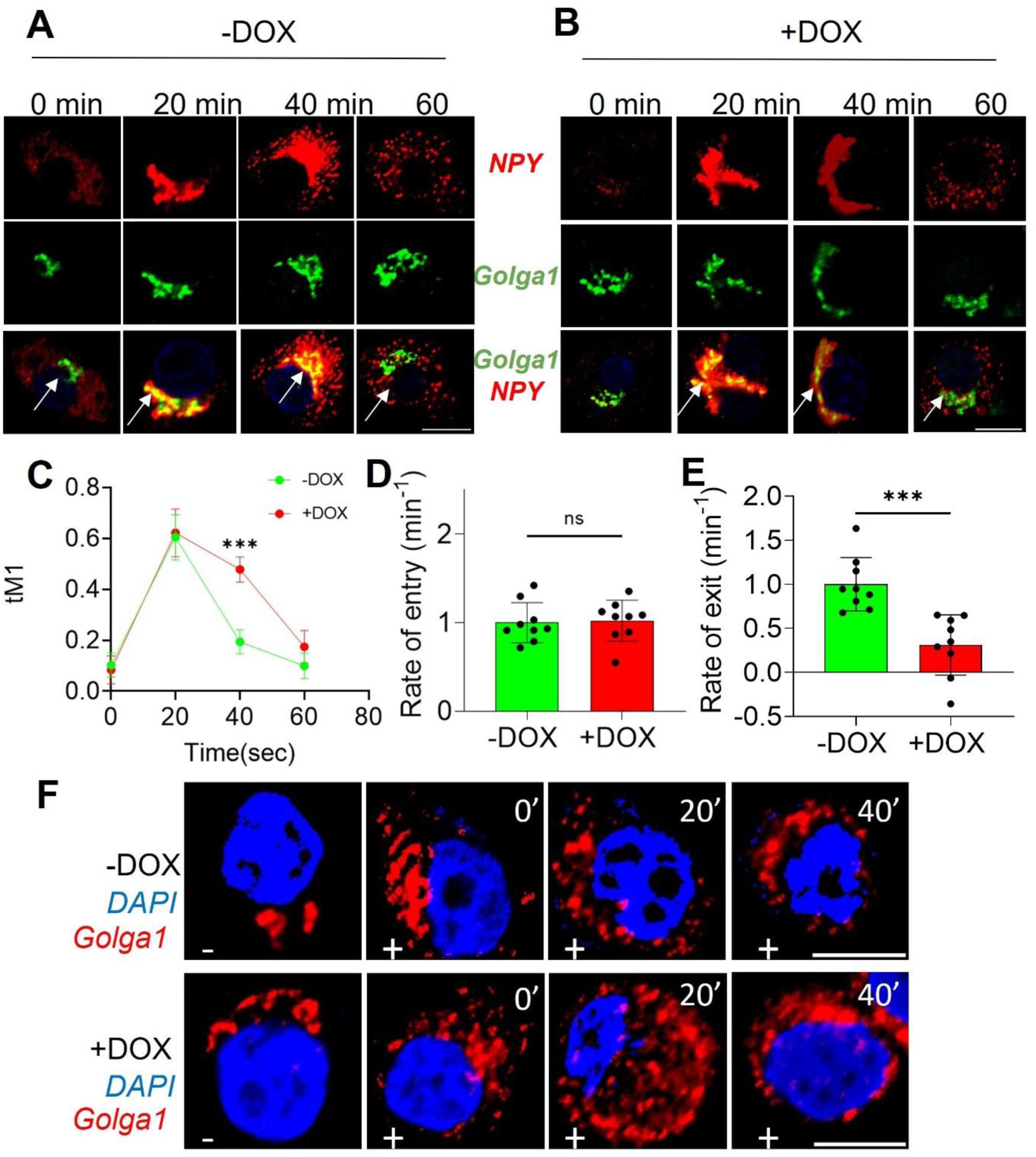
AP-3 affects the dynamic organization of the trans-Golgi network and its subsequent sorting of DCV cargoes. (a-b) Representative confocal microscope fluorescent images for mock-treated and shRNA2 AP-3 depleted cells, where Golga1 is green and NPY is red. Scale bar: 10µm. (c) Graphs showing tM1 (Mander’s coefficient 1) of NPY-mCherry and Golga1 for mock-treated (Green) and shRNA1 AP-3 depleted cells (Red). (d) Graphs for the Rate of entry of Cargo at the Golgi per minute. (e) Graphs show a reduction in Cargo’s exit rate from the Golgi per minute due to AP-3 KD. (N=2 independent experiment, n=10 cells each) (f) Representative confocal images for mock-treated and shRNA1 AP-3 depleted cells treated with Brefeldin A (dis-assembles Golgi) showing a defect in the Golgi reassembly upon AP-3 KD. Scale bar: 10µm. (*P<0.05, **P<0.01, ***P<0.001; n.s. statistically non-significant).

**Fig. S4.**
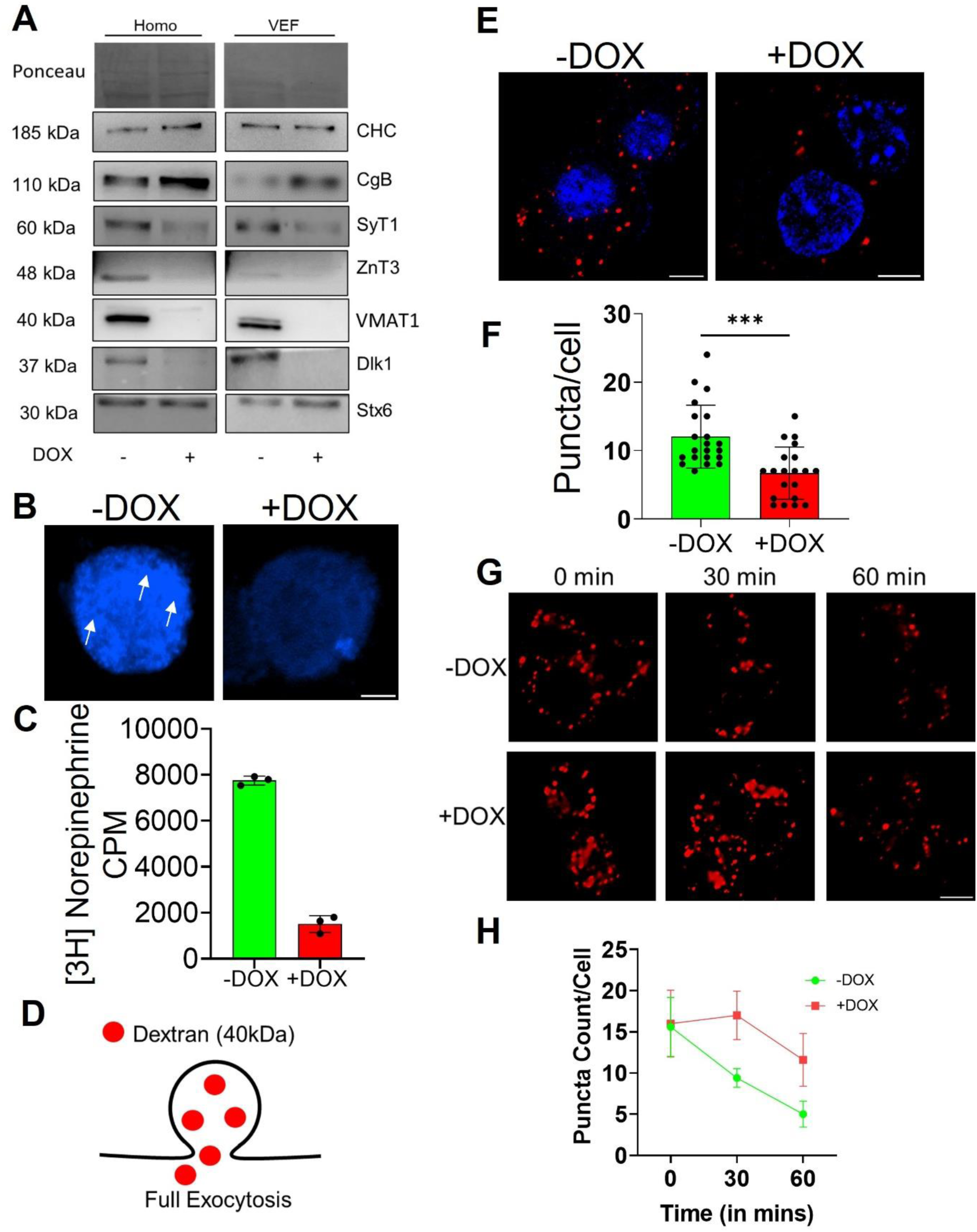
AP-3 plays a selective role in sorting membrane proteins and also affects the trafficking of vesicles post-Golgi in PC12 cells. (a) WB validates the abundance in the cell homogenate and VEFs for the proteins deficient in MS for Mock vs shRNA2 AP-3 depleted cells. (b) Representative fluorescent images of the mock-treated and shRNA2 AP-3 depleted cells showed reduced Zinquin ethyl ester dye uptake. Scale bar: 5µm. (c) The graph depicts approximately five-fold more uptake of Nor-epinephrine in the mock-treated cells after adding up the counts in both cells and the medium. (***Indicates statistical significance at the *p* < 0.001 level assessed by two-tailed Student’s *t*-test (three independent experiments) (d) Schematic for the full exocytotic events. (e-f) Representative images and quantification for Dextran 40kDa show impaired full exocytosis as the puncta per cell reduces in shRNA2 AP-3 depleted cells (Red) w.r.t Mock treated cells (Green). (N=3, n=20 cells each). (g-h) Representative confocal images and quantification for the Endolysosomal experiment using Dextran 10kDa shows a defect in the clearance of the dextran puncta (No. of Puncta/cell) in both the 30 mins and 60 mins in case of shRNA2 AP-3 depleted cells. (N=3, n=20 cells, *P<0.05, **P<0.01, ***P<0.001; n.s. statistically non-significant).

**Fig. S5a-5c:**
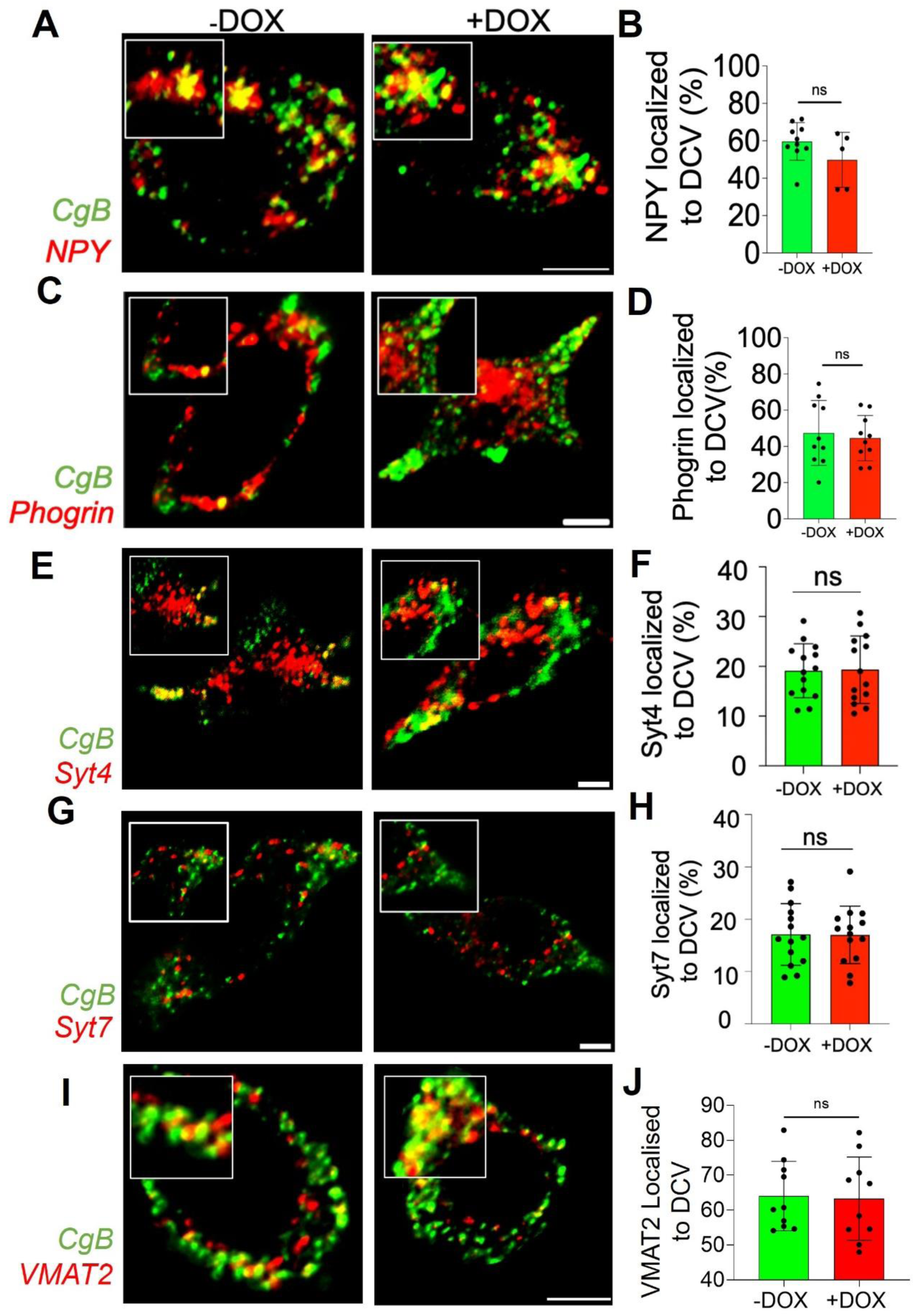
AP-3 preferentially traffics certain membrane proteins to DCVs, which gets missorted to the lysosomes upon depleting AP-3 in PC12 cells. S5a. Representative confocal images for the colocalization of other membrane proteins which are unaffected due to AP-3 depletion. (a-j) NPY, Phogrin, Syt4, and Syt7 (Red) colocalization with DCV marker CgB (Green) does not change significantly in mock-treated and shRNA1 AP-3 depleted cells. For NPY-CgB (n=10 cells for Mock and n=8 cells for shRNA1 AP-3 KD cells). For Phogrin-CgB (n=10 cells each), For Syt4-CgB, Syt7-CgB (n=14 cells each). (*P<0.05, **P<0.01, ***P<0.001; n.s. statistically non-significant).

**S5b.**
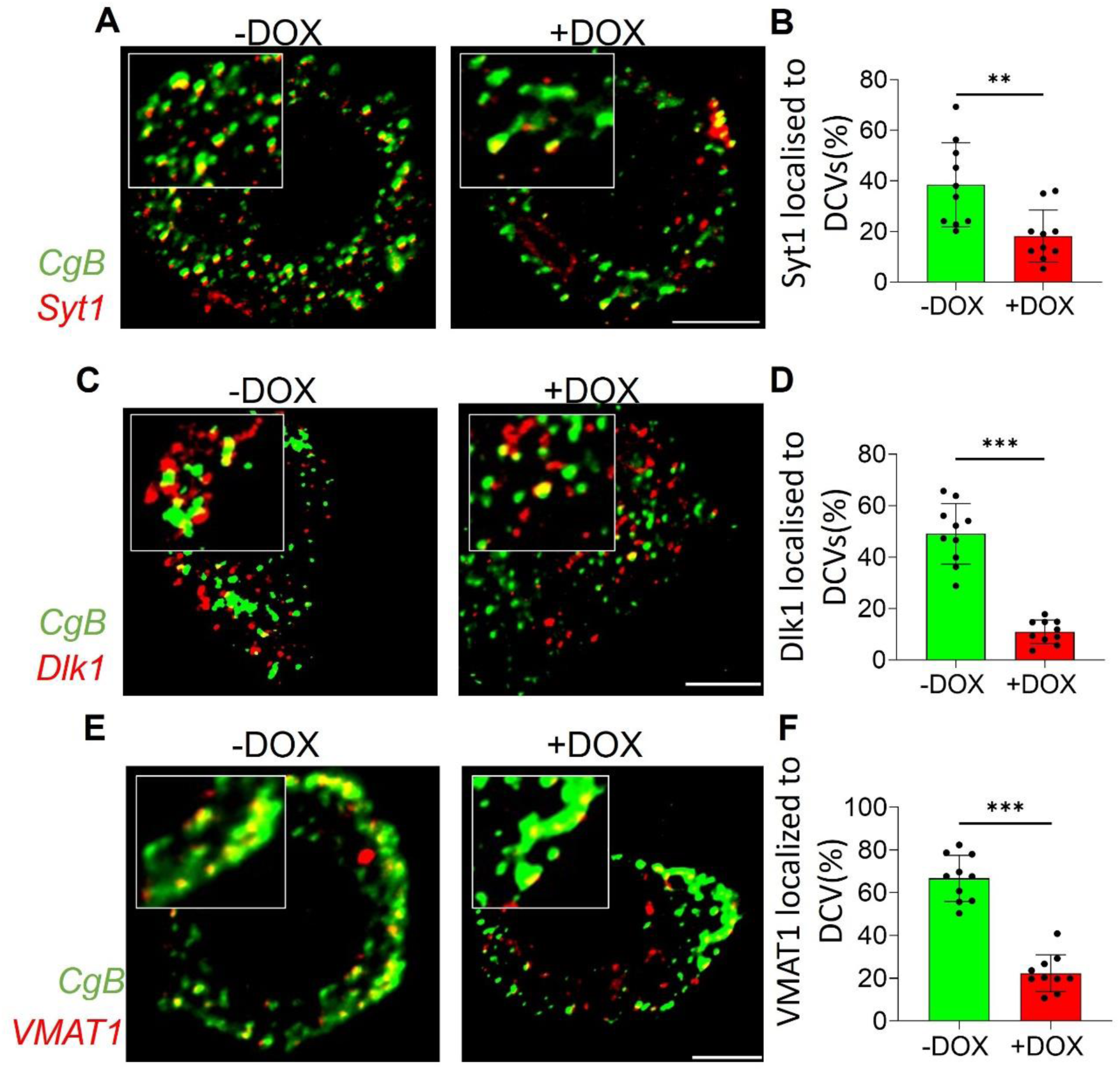
Representative confocal images showing colocalization of specific proteins like Syt1, Dlk1 and VMAT1 with DCV marker CgB in mock vs shRNA2 AP-3 depleted cells. (a-f) Colocalization of Syt1, Dlk1 and VMAT1 (Red) with DCV(Green) reduces in AP-3 knockdown. (n=10 cells or 3 independent experiments for both mock treated and shRNA1 AP-3 depleted cells, *P<0.05, **P<0.01, ***P<0.001; n.s. statistically non-significant).

**S5c.**
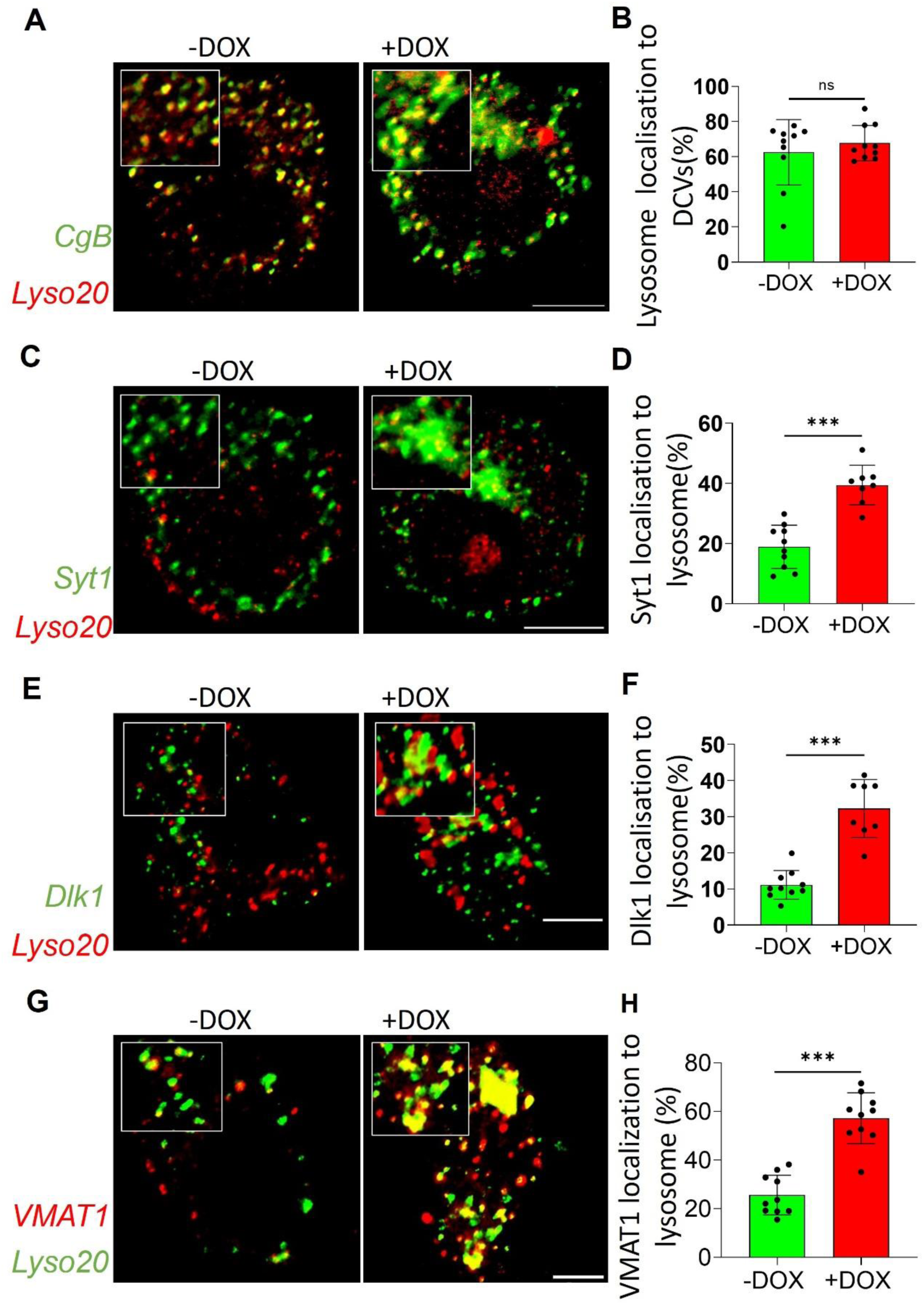
Representative confocal images showing colocalization of Syt1, Dlk1, VMAT1 and CgB with Lysosomal marker LAMP1 in mock vs shRNA2 AP-3 depleted cells. (a-b) Colocalization of DCV marker CgB (Green) with Lysosomal marker Lyso20 or LAMP1 (Red) shows no significant change. (c-j) Colocalization of Syt1, Dlk1 and VMAT1 (Green) with lysosomal marker Lyso20 or LAMP1 (Red) increases AP-3 knockdown, indicating mis-sorting to the lysosomes. (n=10 cells or 3 independent experiments for both mock treated and shRNA1 AP-3 depleted cells, *P<0.05, **P<0.01, ***P<0.001; n.s. statistically non-significant).

**Fig S6.**
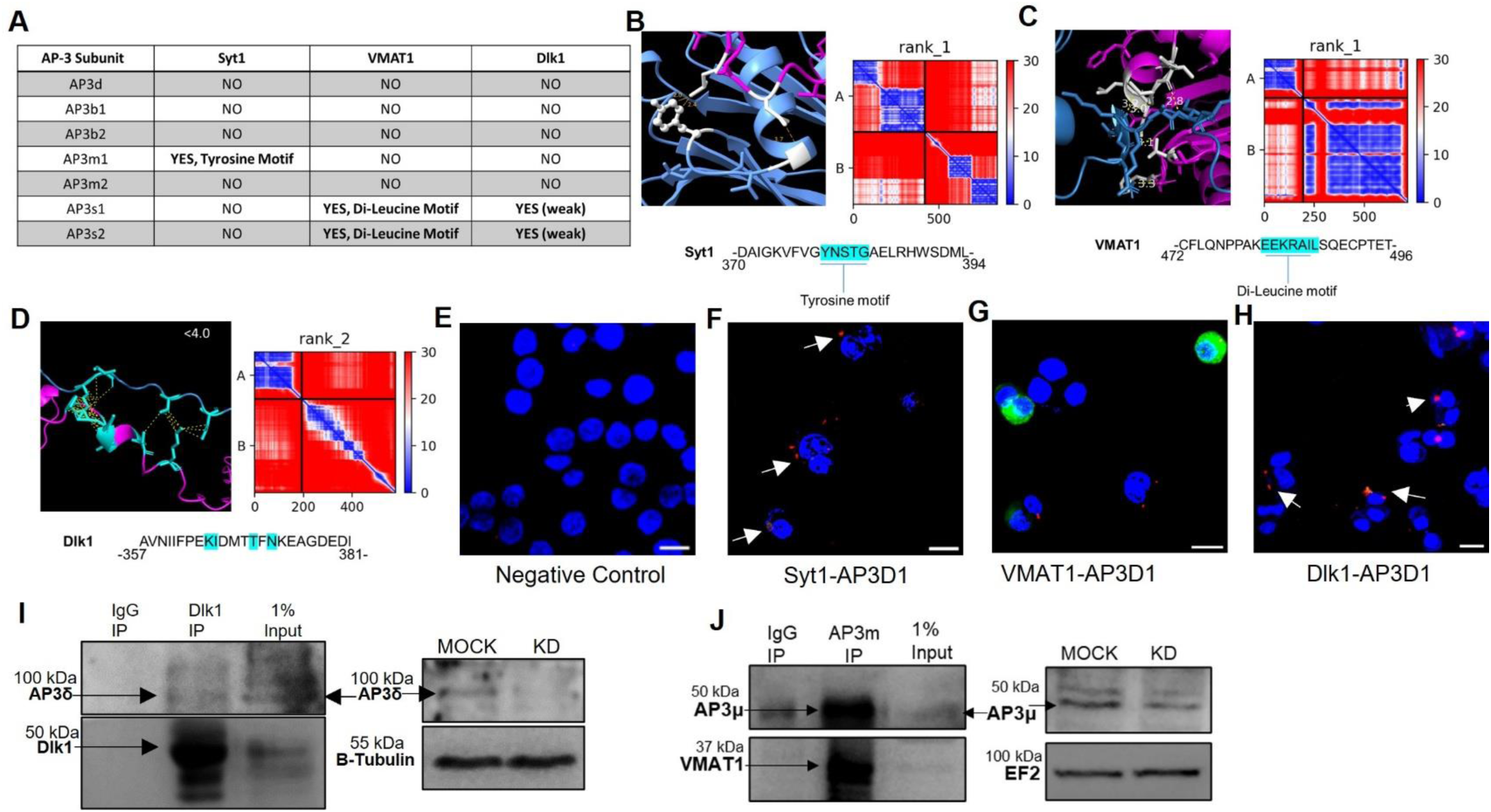
Protein-protein interactions of AP-3 complex with Specific cargoes. (a) Colabfold generated protein-protein interactions of AP-3 subunits with Syt1, Dlk1, and VMAT1. The table depicts the interactions along with the motif involved. (b) AP-3 subunit m1 interacts with Syt1 using a tyrosine motif, and the respective figure shows the interaction site and the PAE plot. (c) AP-3 subunit s1 interacts with VMAT1 using a Di-leucine motif, and the respective figure shows the interaction site and the PAE plot. (d) AP-3 subunit s2 interacts with Dlk1 using an acidic-rich cluster motif, and the respective figure shows the interaction site and the PAE plot. (e-h) Representative confocal images of proximity ligation assay in PC12 cells for showing proximity between Syt1(f), VMAT1(g), and Dlk1(h) with AP3δ, where the red dots indicate the interactions and (e) is the negative control (No antibody). (i) Dlk1 co-immunoprecipitation blots in wild type PC12 cells showing a pulldown of the AP-3delta subunit and Dlk1. The blots on the right are represented to indicate the bands specific to AP-3delta subunit in the cell lysates of mock vs the AP-3 depleted cells. (j) AP-3mu subunit co-immunoprecipitation blots in wild type PC12 cells showing a pulldown of the AP-3mu subunit and VMAT1. The blots on the right are represented to indicate the bands specific to AP-3mu subunit in the cell lysates of mock vs the AP-3 depleted cells.

**Figure S7.**
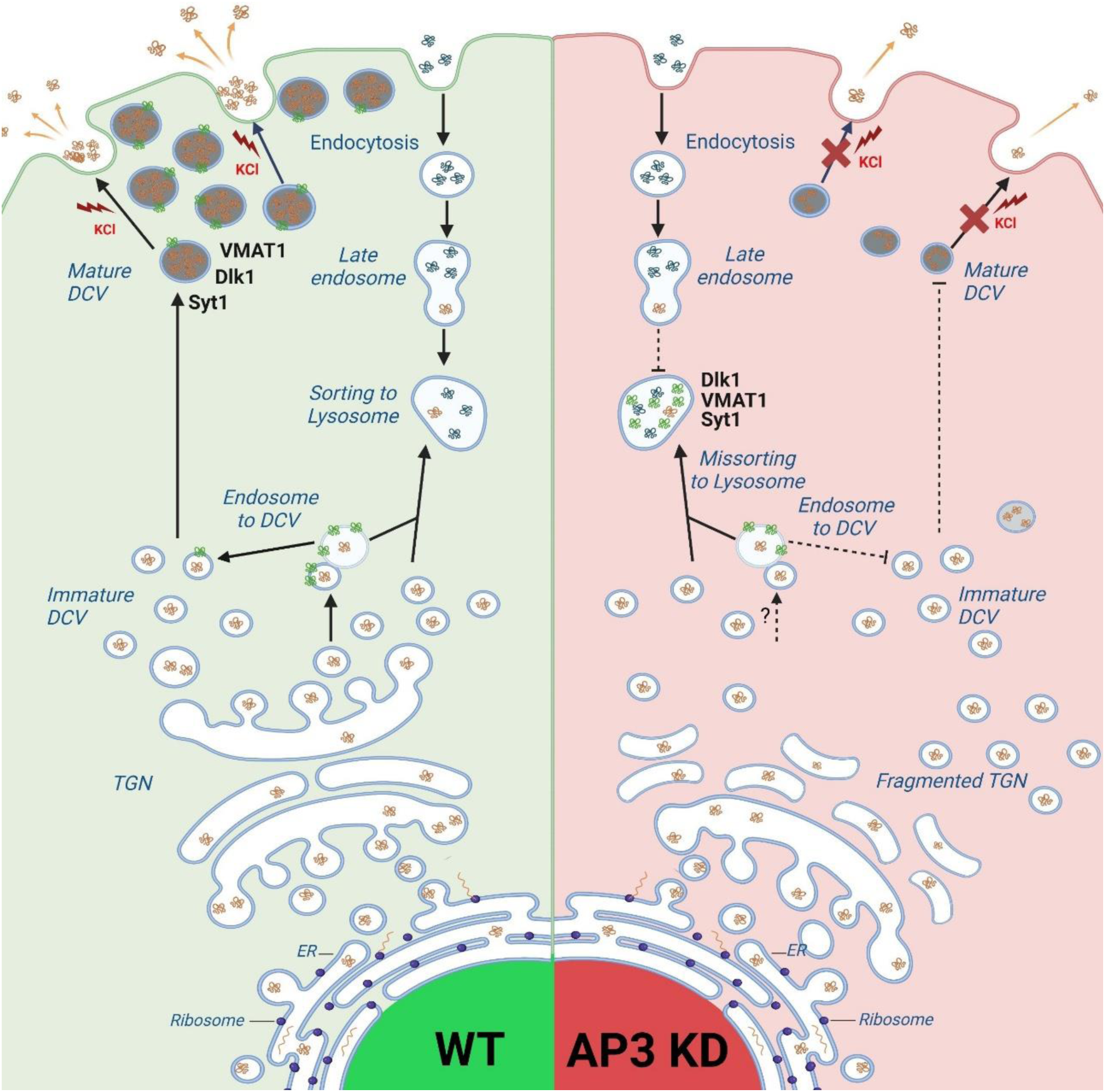
Plausible mechanisms of AP-3 affecting DCV function. The presence of AP-3 at both the TGN and the endosomes gives rise to two plausible mechanisms of cargo sorting. At the TGN, AP-3, along with clathrin, helps form specific sorting sites where the DCV cargo is loaded into the immature DCVs. These immature DCVs then undergo homotypic fusion and mature, moving farther away from the TGN and getting docked at the plasma membrane for stimulus-coupled release. Alternatively, AP-3 present at the endosomes may be responsible for loading DCV cargo into the immature vesicles, thus making the DCVs more condensed and helping the maturation. AP-3 is also responsible for the budding of SLVs at the endosomes, the recycling of exocytotic vesicles and clearance via the Endolysosomal pathway. This figure shows the vesicle trafficking in WT PC12 cells compared to AP-3 KD cells. The absence of AP-3 is responsible for inhibiting the maturation of DCVs and may lead to stunted transport of specific membrane proteins and Cargo to the DCVs. Loss of AP-3 leads to the mis-sorting of membrane proteins to the lysosomes. Thus, it affects the regulated exocytosis of DCV at multiple checkpoints.

**Video S1-S6. Related to Figures 3,S1 and 4**

**Video S1-AP3 shRNA1 -DOX NPY pHtomato video**

The video shows NPY-pHtomato live cell imaging with KCl stimulation added at 2 seconds in mock or -DOX AP3 shRNA1 cells. It runs at 30 frames per second.

**Video S2-AP3 shRNA1 +DOX NPY pHtomato video**

The video shows NPY-pHtomato live cell imaging with KCl stimulation added at 2 seconds in +DOX AP3 shRNA1 Knockdown cells. It runs at 30 frames per second.

**Video S3-AP3 shRNA2 -DOX NPY pHtomato video**

The video shows NPY-pHtomato live cell imaging with KCl stimulation added at 2 seconds in mock or -DOX AP3 shRNA2 cells. It runs at 30 frames per second.

**Video S4-AP3 shRNA2 +DOX NPY pHtomato video**

The video shows NPY-pHtomato live cell imaging with KCl stimulation added at 2 seconds in +DOX AP3 shRNA2 Knockdown cells. It runs at 30 frames per second.

**Video S5-AP3 shRNA1 -DOX Synaptophysin pHtomato video**

The video shows Synaptophysin-pHtomato live cell imaging with KCl stimulation added at 2 seconds in mock or -DOX AP3 shRNA1 cells. It runs at 20 frames per second.

**Video S6-AP3 shRNA1 +DOX Synaptophysin pHtomato video**

The video shows Synaptophysin-pHtomato live cell imaging with KCl stimulation added at 2 seconds in +DOX AP3 shRNA1 Knockdown cells. It runs at 20 frames per second.

